# Non-targeting control for MISSION^®^ shRNA library silences *SNRPD3* leading to cell death or permanent growth arrest

**DOI:** 10.1101/2020.11.25.398354

**Authors:** Maria Czarnek, Katarzyna Sarad, Agnieszka Karaś, Jakub Kochan, Joanna Bereta

## Abstract

In parallel with the expansion of RNA interference techniques evidence has been accumulating that RNAi analyses may be seriously biased due to off-target effects of gene-specific shRNAs. Our work points to another possible source of misinterpretations of shRNA-based data – off-target effects of non-targeting shRNA. We found that one such control for the MISSION^®^ library (commercialized TRC library), SHC016, is cytotoxic. Using a lentiviral vector with inducible expression of SHC016 we proved that this shRNA induces apoptosis in murine cells and, depending on p53 status, senescence or mitotic catastrophe in human tumor cell lines. We identified SNRPD3, a core spliceosomal protein, as a major SHC016 target in several cell lines and confirmed in A549 and U251 cell lines that CRISPRi-knockdown of *SNRPD3* mimics the effects of SHC016 expression. Our finding disqualifies non-targeting SHC016 shRNA and adds a new premise to the discussion about the sources of uncertainty of RNAi results.

## Introduction

RNA interference (RNAi) is one of the leading methods to study gene functions via loss of function approach. The technique is based on the natural mechanisms of post-transcriptional gene silencing by short RNA species such as miRNA derived from endogenous precursors and siRNA derived from exogenous long dsRNA of viral origin (Shi, 2003).

In mammalian cells primary microRNAs (pri-miRNAs) are processed in the nucleus by Microprocessor complex to produce hairpin precursor miRNAs (pre-miRNAs) which are further processed by the cytosolic endonuclease Dicer to miRNA duplexes with two-nucleotide overhangs on 3’ end of each RNA strand. Argonaute protein is loaded with one strand of the miRNA duplex to create RNA-Induced Silencing Complex (RISC), which interacts with a target mRNA leading to inhibition of its translation and/or degradation (Gebert & MacRae, 2019).

Initially miRNA-like siRNA duplexes, not requiring endonucleolytic processing were designed and introduced into cells via transfection to silence the genes of interest. However, inefficient transfection of many cell lines as well as transience of siRNA activity led to the development of short hairpin RNA (shRNA) expression vectors including retro- and lentiviral vectors (Paddison, Caudy, Sachidanandam, & Hannon, 2004). The expression cassette coding for a given shRNA is stably integrated into DNA of transduced cells and the transcribed shRNA, which mimics pre-miRNA, is processed by Dicer to an siRNA duplex (Sheng, Flood, & Xie, 2020).

The need for unified research tools has triggered the development of shRNA libraries which in the same shRNA backbone contain sequences that silence individual genes. The engineering of shRNA libraries allowed also for the development of high-throughput methods employing arrayed or pooled RNAi screens to identify proteins involved in different cellular processes or novel specific therapeutic targets (Berns et al., 2004; Chang, Elledge, & Hannon, 2006; Moffat et al., 2006; Paddison, Silva, et al., 2004).

One of the widely used shRNA libraries is the lentiviral TRC library developed by The RNAi Consortium at the Broad Institute and commercially available through Merck (Darmstadt, Germany; previously Sigma-Aldrich, St. Louis, MO) under the trade name MISSION^®^. The library contains shRNAs targeting ~15 thousand human and ~15 thousand mouse transcripts; each transcript is covered by an average of 5 shRNAs. The library was generated as pLKO.1 lentiviral vectors containing puromycin-resistance gene and shRNA sequences flanked by Pol III-recognized elements, human U6 promoter and T-stretch termination signal (Moffat et al., 2006).

Each RNAi experiment, regardless of whether it is a high throughput screening or analysis of individual transcripts, requires proper controls. *“Empty vector control”* does not contain any shRNA sequence and allows to evaluate the effects of the transduction procedure and of lentiviral vector elements (other than shRNA) on studied cells. The second type of control, *“non-targeting shRNA”,* contains an shRNA-coding sequence that does not target any gene of the studied species. After the transduction of the cells with this control lentiviral vector, shRNA is produced and influences the miRNA processing machinery and possibly other cellular processes in a way corresponding to the unspecific effects triggered by gene-targeting shRNAs. This control should be the reference for all experimental results.

MISSION^®^ library provides two non-targeting shRNA controls. The first one, designated SHC002, does not target any mammalian transcripts. However, it targets turboGFP transcript and therefore is not recommended for tGFP-expressing cells. According to the supplier information, the second non-targeting shRNA control, designated SHC016, has been bioinformatically determined not to target any transcript in any species (Merck, 2020).

We have used this control, SHC016, in our research and observed that it had detrimental effects on transduced cells. We ruled out the possibility that the high rate of cell death was due to puromycin effect on the cells that remained non-transduced. To do so, we generated vectors with tetracycline/doxycycline-inducible expression of control shRNAs, either SHC002 or SHC016, which enabled us to separate in time the process of transduction/selection from the process of shRNA expression. Using these vectors, Tet-on-SHC002 and Tet-on-SHC016, we proved cytotoxicity of SHC016 shRNA. We also demonstrated that SHC016 induces different cell death pathways in murine and human cell lines and elucidated its mechanism of action.

## Results

We suspected that the MISSION^®^ non-targeting control shRNA, SHC016, is deleterious to transduced cells. To verify this notion we compared the effects of doxycycline-induced two nontargeting shRNAs, SHC002 and SHC016. MC38CEA cells were transduced with the same amounts of either Tet-on-SHC002 or Tet-on-SHC016 and the cells with incorporated transgenes were selected with puromycin. Then the cells were cultured for five days with the expression of the nontargeting shRNAs turned on for the last 4, 3, or 2 days. Unlike SHC002, SHC016 had a huge impact on transduced MC38CEA. The viability of the cells dropped to ~30% on the fourth day after inducing SHC016 expression *(Figure 1A).* The increased exposure of phosphatidylserine on the external leaflet of the plasma membrane detected by annexin V binding *(Figure 1B and Figure 1— figure supplement 1A)* and enhanced activity of executioner caspases 3 and 7 *(Figure 1C)* indicated that the expression of SHC016 induced apoptosis in MC38CEA cells. This process was accompanied by increasing number of the cells in G2/M and decreasing number of the cells in G1 phase of the cell cycle, which indicated the cell cycle arrest at the G2/M phase *(Figure 1D and Figure 1—figure supplement 1B).*

**Figure 1.**
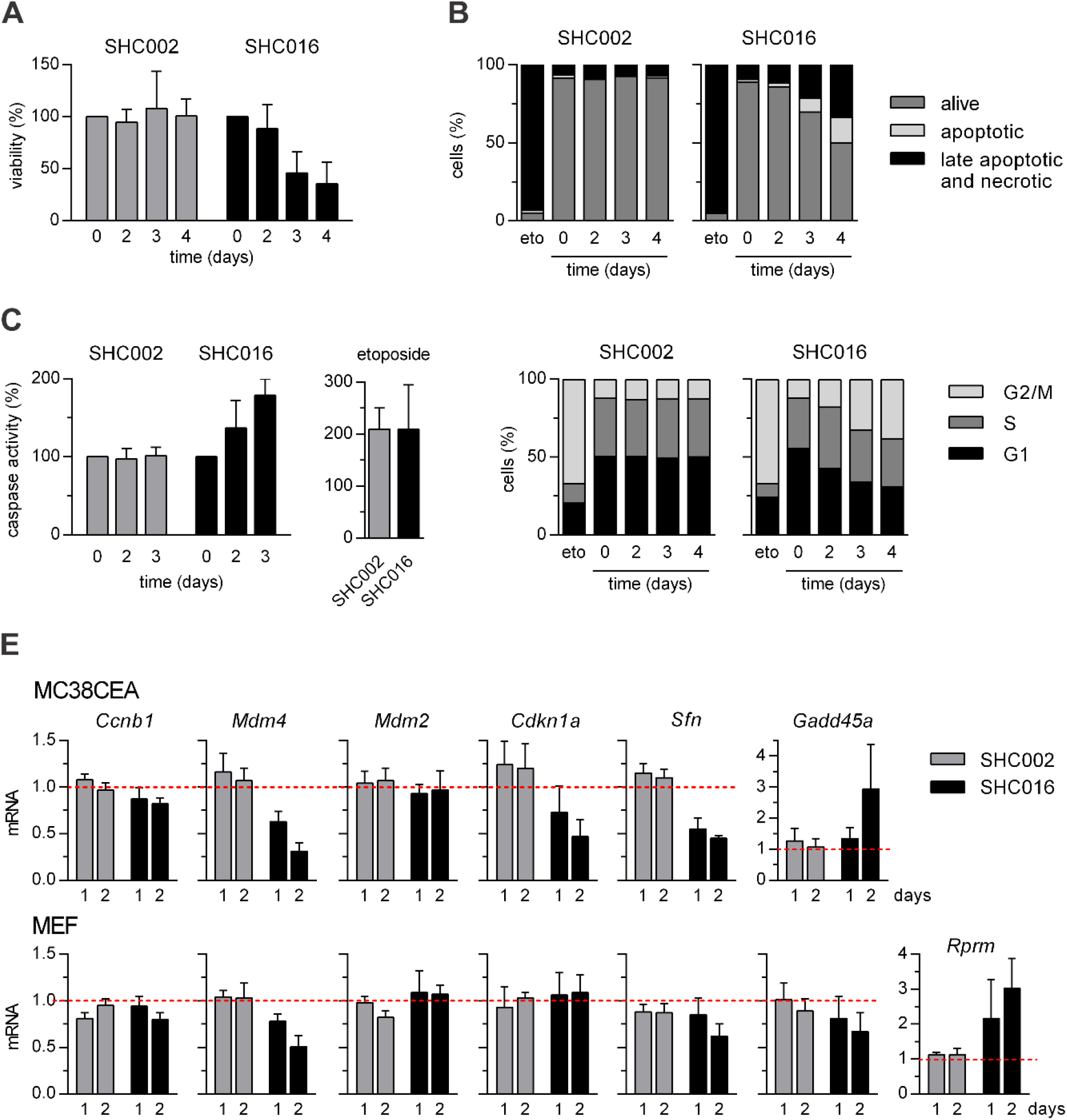
Expression of SHC016 in MC38CEA results in apoptosis and cell cycle arrest at the G2/M phase. The expression of non-targeting shRNA SHC002 or SHC016 was induced by doxycycline (100 ng/ml) for the last 4, 3, or 2 days of the 5-day culture. (A) Cell viability assessed by MTT assay. Absorbance values of the cells without induced transgene expression were taken as 100%. (B) Analysis of death via apoptosis assessed by annexin V/PI staining and flow cytometry analysis; alive cells – annexin V^−^/PI^−^; apoptotic – annexin V^+^/PI^−^; late apoptotic and necrotic – annexin V^+^/PI^+^ and annexin V^−^/PI^+^ (the latter usually did not exceed 3% of the cells expressing either shRNA). (C) Activity of caspases 3 and 7 measured via chemiluminescence assay. Chemiluminescence values of the cells without induced transgene expression were taken as 100%. (D) Flow cytometry analysis of the cell cycle. (B-D) The cells treated for the last 2 days of experiment with etoposide (2 μM) were used as a positive control. (B, D) Different representations of the data containing error bars can be found in *Figure 1—figure supplement 1A, B.* (E) RT-qPCR analysis of mRNA levels of p53-target genes in MC38CEA and MEF cells. (A-E) Data are shown as mean values (MV) from 3 independent experiments. Error bars represent standard deviation (SD).

In another murine cell model, murine embryonal fibroblast cell line (MEF), both the time of appearance and the magnitude of SHC016-mediated effects on cell viability and apoptosis resembled those observed for MC38CEA *(Figure 1—figure supplement 2).* However, unlike for MC38CEA, the impact of SHC016 on the cell cycle of MEF was weak and detectable only in the fourth day of its expression, but similarly to MC38CEA, increased number of cells in G2/M phase was observed. We excluded the so-called interferon response as a cause of SHC016-mediated effects because of a lack of increased expression of interferon-inducible genes, *Oas1b*, *Ifit1*, and *Pkr (Figure 1—-figure supplement 3).*

Because the accumulation of p53 is believed to be the major trigger of G2/M arrest we analyzed, in both MC38CEA and MEF, the influence of SHC002 and SHC016 on the expression of p53-target genes including *Mdm2*, apoptosis inducers *Bbc3* (also known as *Puma*) and *Pmaip1* (*Noxa*) as well as those involved in inhibition of G2/M transition such as *Cdkn1a* (coding for p21), *Gadd45a, Sfn* (stratifin, 14-3-3σ), and *Rprm* (reprimo) (Fischer, 2017; Ohki et al., 2000). We added to the analysis *Ccnb1*, which encodes mitotic cyclin (cyclin B1) and *Mdm4*, whose product, apart from MDM2, binds p53 and inhibits its transcriptional activity (Karni-Schmidt, Lokshin, & Prives, 2016). Although, in both MC38CEA and MEF notable changes in gene expression were observed only in the cells with induced expression of SHC016, but not of SHC002, the results were not consistent (*Figure 1E*). The expression of *Ccnb1* was only slightly diminished in either cell line. The decrease in *Mdm4* expression observed in both cell lines did not turn on a typical p53 response because the levels of p53-target genes were mostly unaffected *(Bbc3, Pmaip1;* data not shown) or even reduced *(Cdkn1a* in MC38CEA and *Sfn* in both lines). The augmented expression of *Gadd45a* was observed only in MC38CEA. The level of *Rprm,* which is undetectable in MC38CEA, was upregulated in SHC016-expressing MEF. The expression of both, *Gadd45a* and *Rprm,* may be upregulated not only by p53 but also by other transcription factors (Amigo et al., 2018; Lal & Gorospe, 2006). GADD45A is believed to affect cell cycle progression via interactions with various players including PCNA, CDK1 and p21. Reprimo inhibits formation of active cyclin B-CDK1 complex. Consequently, the expression of SHC016 induced distinct sets of changes in the expression of genes related to cell cycle control in the two mouse cell lines. Although the changes may affect G2/M transition in both MC38CEA and MEF cells, they do not imply the mechanism of SHC016 action.

We further analyzed the effect of the non-targeting sequences SHC002 and SHC016 on five human tumor cell lines, A549, U251, HeLa, PC3, and MCF7 to examine whether the deleterious effects of SHC016 are limited to cells of mouse origin, or are a more common phenomenon involving also human cells. In contrast to SHC002, SHC016 had a potent impact on viability of all studied cell lines *(Figure 2A* and *Figure 2—figure supplement 1*).

**Figure 2.**
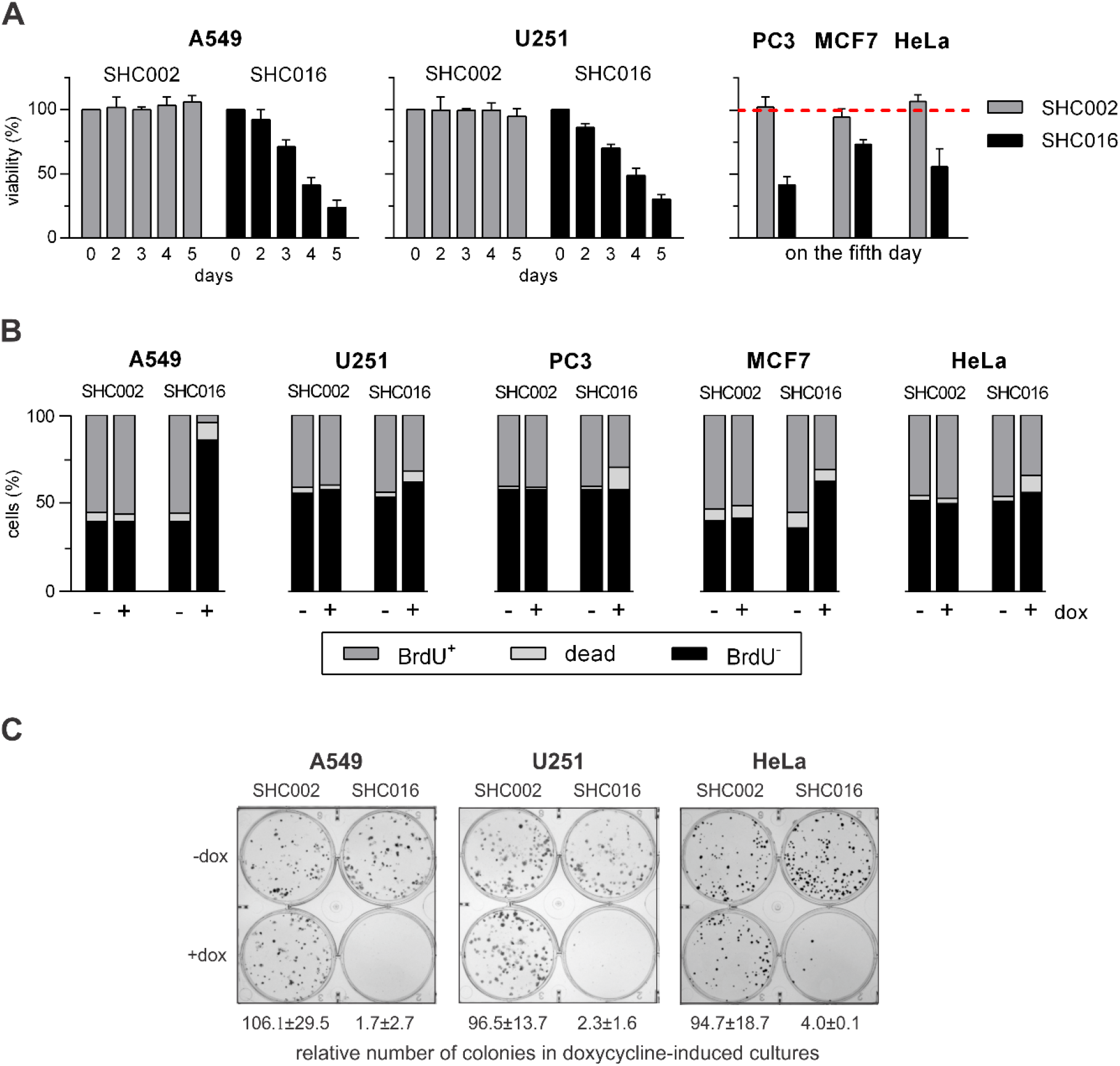
Expression of SHC016 in human cell lines results in inhibition of cell proliferation and/or viability. (A) Cell viability assessed by MTT assay. The expression of non-targeting shRNA SHC002 or SHC016 was induced by doxycycline (dox, 100 ng/ml) for the last 5, 4, 3, or 2 days of the 6-day culture. The measurements taken each day (A549, U251) or on the fifth day of dox treatment (PC3, MCF7, HeLa) are shown. Full data for PC3, MCF7, and HeLa are available in *Figure 2—figure supplement 1.* Absorbance values of the cells without induction of shRNA expression were taken as 100%. (B) DNA synthesis and cell viability assessed via double BrdU/eFluor 520 viability staining performed three (A549) or five days (U251, PC3, MCF7, HeLa) after inducing SHC002-or SHC016 expression. BrdU was added to the medium for the last 6 h of culture; dead – all eFluor 520^+^ cells, BrdU^+^ – eFluor 520^−^/BrdU^+^ cells, BrdU^−^ – eFluor 520^−^/ BrdU^−^ cells. Different representations of the data containing error bars can be found in *Figure 2—figure supplement 4.* (A, B) Data are shown as MV from 3 independent experiments. Error bars in (A) represent SD. (C) Analysis of clonal growth capacity of A549, U251, and HeLa cells by colony formation assay.

We excluded the possibility that the negative effects of SHC016 was a matter of a high transgene levels because qPCR analysis indicated that the levels of a pLKO cassette coding for SHC016 were comparable to or lower than that coding for SHC002 in all analyzed cell lines *(Figure 2—figure supplement 2).* Also, the interferon response did not occur either in A549 or U251 cells *(Figure 2—figure supplement 3*).

Unlike for murine cells, apoptosis, as evaluated by annexin V/PI test, seemed not to play a primary role in SHC016-mediated effects in human cells (data not shown). The cell cycle analysis did not bring conclusive results. Therefore, to answer the question whether SHC016 sequence affects proliferation or survival of human cells we performed double proliferation/viability test, in which incorporation of BrdU indicating ongoing DNA synthesis was analyzed in parallel with staining of dead cells after inducing non-targeting shRNAs expression (*Figure 2B* and *Figure 2—figure supplement 4)* Only in A549 cells the incorporation of BrdU was almost completely inhibited upon induction of SHC016 expression. For U251, PC3, and HeLa cell lines the percentage of cells that had ceased to synthetize DNA, did not increase or increased only slightly, while in MCF7 the decline in BrdU incorporation was notable but much less spectacular than in A549. Interestingly, the type of response correlates with p53 status of the cell lines. Only A549 is known to possess fully active p53. Both U251 and PC3 express only mutated variants of p53, unable to activate transcription of p53-target genes and although transcription of wild type *TP53* occurs in HeLa and MCF7 cell lines, the activity of p53 protein is reduced (Leroy et al., 2014). In HeLa cells papilloma virus protease E6 degrades p53 to the level undetectable in western blotting and in MCF7 cells *MDM4* amplification impairs p53 functions (Leroy et al., 2014). However, independently on p53 status, the cells with induced SHC016 expression underwent irreversible growth arrest; they lost capacity to produce colonies, which we demonstrated in a colony formation assay performed for A549, U251 and HeLa cell lines *(Figure 2C).*

Pictures of culture plates with stained clones were taken 7–10 days after plating the cells. Expression of SHC002 or SHC016 was induced with doxycycline 24 h after seeding the cells. The number of colonies in the corresponding, uninduced cultures were taken as 100%. Data are shown as MV ± SD from 3 (U251 and HeLa) or 4 (A549) independent experiments.

We presumed that SHC016 exerted its deleterious effects on the cells via different mechanisms, one of which involved activation of p53 and occurred in A549 cells. To confirm this notion and further elucidate the possible mechanism of SHC016-induced effects in A549 cells we analyzed, using RT-qPCR, the potential changes in the expression of genes (*i*) activated by p53, including these coding for cell cycle inhibitors and mediators of apoptosis *(CDKN1A, MDM2, BBC3, PMAIP1, GADD45A, SFN), (ii)* coding for central players in cell cycle progression including *CDK2, CDK1, CCNB1, AURKA, PLK1,* components of chromosomal passenger complex, abbreviated as CPC *(AURKB, BIRC5, CDCA8)* and (*iii*) coding for proteins of cell cycle checkpoints (*CHEK1*, *CHEK2*, *BUB1*, *BUB1B*, *BUB3*, *MAD2L1*). Turning on the expression of SHC016, but not that of SHC002, resulted in the stimulation of p53-regulated genes: *CDKN1A*, *MDM2*, and *BBC3* already 24 h after addition of doxycycline to the culture medium. The effect was further enhanced after 48 h (*Figure 3A*). The strongest stimulation was observed for *CDKN1A*, which codes for p21, a universal inhibitor of cyclin-dependent kinases. Additionally, p21 activates the formation of DREAM complex, which plays the role of a transcriptional repressor of genes involved in cell cycle progression including all genes from our group (*ii*) and (*iii*) (Engeland, 2018). This explains diminished levels of the majority of analyzed transcripts 48 h after inducing expression of SHC016 *(Figure 3—figure supplement 1*) and halting of A549 proliferation.

**Figure 3.**
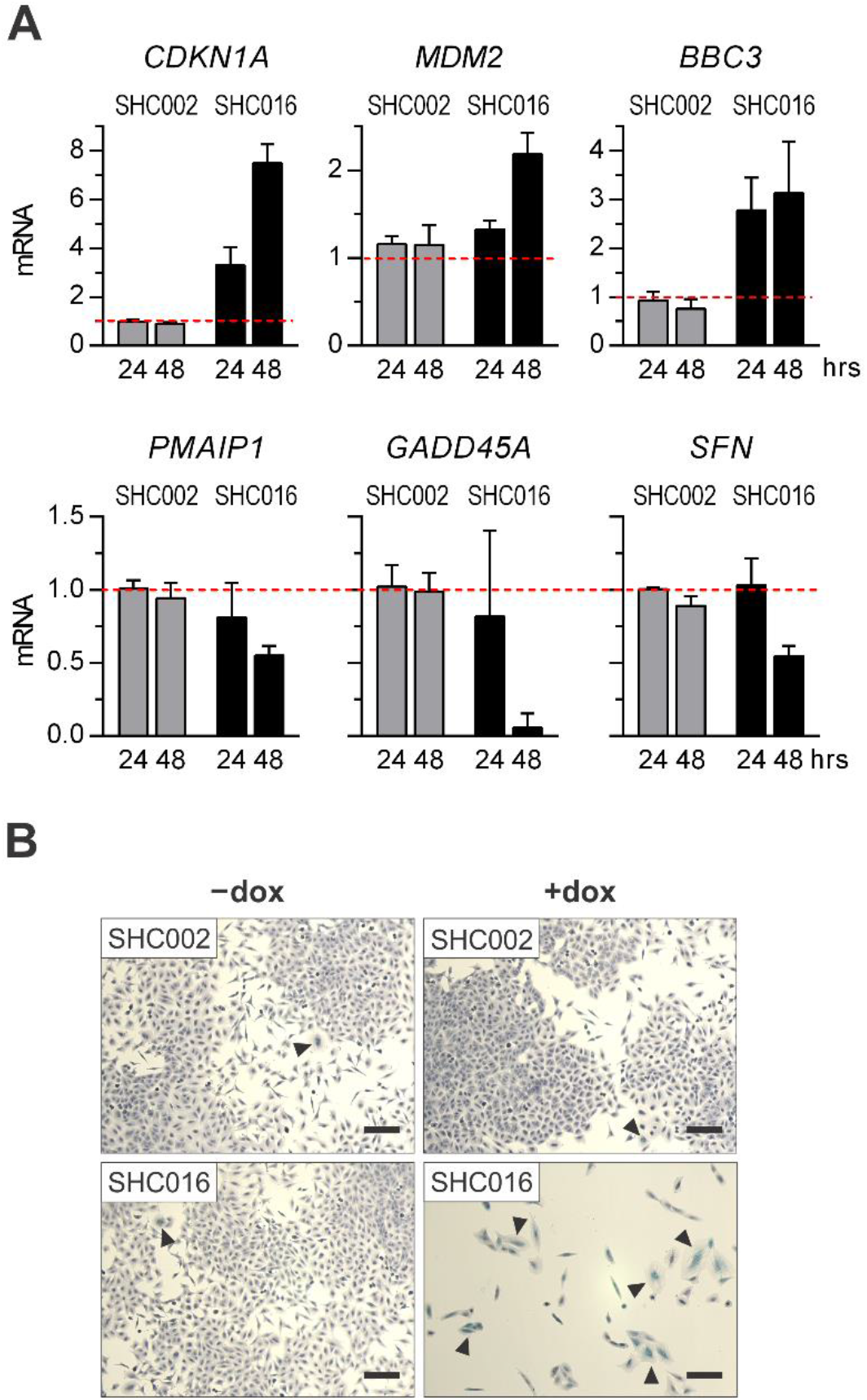
Expression of SHC016 in A549 cells initiates cell senescence. (A) RT-qPCR analysis of mRNA levels of p53-target genes. The relative levels of the transcripts in uninduced cells were taken as 1. MV ± SD from 3 independent experiments are shown. (B) Representative images of A549 cells stained for SA-β-gal activity and counterstained with hematoxylin. The cells were left untreated or the expression of SHC002 or SHC016 was induced with doxycycline (dox) five days prior to staining. Exemplary SA-β-gal-positive cells, very rare among untreated- and SHC002-expressing cells but common among SHC016-expressing cells are indicated by arrowheads. The scale bar represents 100 μm. Three independent experiments were performed.

Additionally, the analysis of *CDKN1A* expression in all studied cell lines showed lack of p53 transcriptional activity in U251, PC3, and MCF7 and only slight activity in HeLa cells *(Figure 3— figure supplement 2*).

The expression of three of potential p53 targets was not increased but rather decreased in A549 cells in response to SHC016: *PMAIP1, GADD45A,* and *SFN (Figure 3A).* The genes could be subject to other regulatory mechanisms. Interestingly, the expression of *Sfn* (mouse ortholog of *SFN)* was diminished in both SHC016-expressing murine cell lines and the expression of *Gadd45a* in one of them, MEF. Thus, it is tempting to speculate that the expression of SHC016 may have an impact on regulation of these transcripts’ levels.

In addition to growth arrest, p21 may trigger a cellular senescence program (Georgakilas, Martin, & Bonner, 2017), in which cells permanently cease proliferation and show significant alterations in morphology but remain metabolically active (Calcinotto et al., 2019; Munoz-Espin & Serrano, 2014; Sikora, Mosieniak, & Sliwinska, 2016). The activity of a key senescence marker, senescence-associated β-galactosidase (SA-β-gal), was observed in majority (59.5% ± 2.13%; mean value ± SD) of A549 cells with induced expression of SHC016. In contrast, the numbers of SA-β-gal-positive cells in the culture of these cells but without SHC016 induction as well as in the cultures of A549 transduced with Tet-on-SHC002 were negligible; they accounted for less than 1% of the cells (*Figure 3B*). The observed enlarged and flattened cell areas as well as their irregular shapes are also characteristic for senescence (*Figure 3B*). We did not observe notable changes in SA-β-gal expression in other studied cell lines; in PC3, HeLa and U251 SA-β-gal remained low despite induction of SHC016, while MCF7 cells express high levels of SA-β-gal independently of transduction with Tet-on vectors and doxycycline induction (data not shown).

In the case of *TP53*-mutant cell lines, U251 and PC3, SHC016 exerts its harmful effects via different mechanism. The images of SHC016-expressing U251 and PC3 cells show the hallmarks of mitotic catastrophe (*Figure 4A and Figure 4—figure supplement 1A, B, D*). Mitotic catastrophe (MC) is defined as a mechanism that senses aberrant mitosis and drive the cells to an irreversible fate (death or senescence) (Mc Gee, 2015). The process may occur when defective checkpoints fail to arrest cell cycle progression in response to critical conditions such as genotoxic stress, a delay in DNA replication, or aberrant spindle formation.

**Figure 4.**
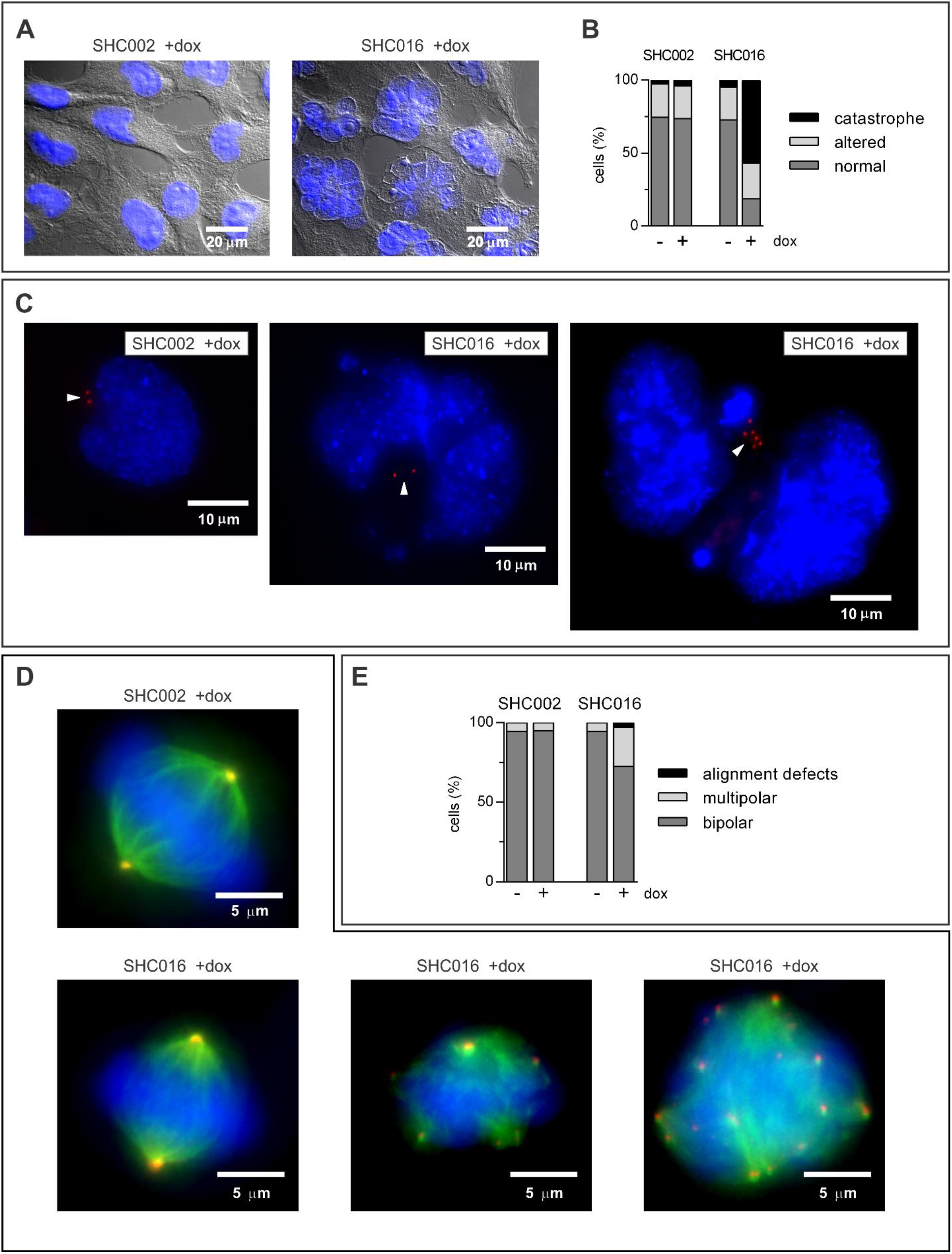
Expression of SHC016 in U251 cells results in mitotic catastrophe. Fluorescence microscopy analyses of nuclear morphology and mitosis in U251 cells. The cells were left untreated or the expression of SHC002 or SHC016 was induced with doxycycline (dox) five days prior to DNA staining. (A) Exemplary merged transmitted light- and fluorescence images representing U251 with regular nuclei typical for SHC002-expressing cells or U251 with catastrophic nuclei prevailing among SHC016-expressing cells. DNA was stained with DAPI (blue). Original images are available in *Figure 4—figure supplement 1A)* More examples are presented in *Figure 4—figure supplement 1B)* (B) Quantitative analysis of nuclear morphology in U251 with uninduced or induced expression of SHC002 or SHC016. Data are from two independent experiments; in each experiment at least 120 cells were analyzed in each experimental group. The population described as ‘altered’ includes cells with significantly enlarged nuclei, nuclei with irregular shapes, and those with micronuclei. Different representation of the results, which include SD values is available in *Figure 4—figure supplement 1C*. (C) Exemplary fluorescence images of interphase nuclei of SHC002- or SHC016-expressing cells. DNA was stained with Hoechst 33342 (blue). Arrowheads point out pairs of centrosomes (left and middle images) or supernumerary centrosomes (right image) visualized by γ-tubulin immunostaining (red). More images are available in *Figure 4—figure supplement 2.* (D) Exemplary fluorescence images of metaphase in U251 cells. Left images present correct number and position of centrosomes and correct formation of the spindle and metaphase plate; remaining images show aberrant number of centrosomes and metaphase plate formation. Spindle microtubules were visualized by β-tubulin immunostaining (green), centrosomes were visualized by γ-tubulin immunostaining (red), and DNA was stained with Hoechst 33342 (blue). (E) Quantitative analysis of correct and aberrant metaphase in U251 cells. The cells were left uninduced or were doxycycline (dox)-induced to express SHC002 or SHC016 two days before analysis. Data are from three independent experiments. Total number of metaphase nuclei analyzed in each experimental group were ~160.

The giant, multinucleated cells, whose formation is typical for MC, made up a significant proportion of SHC016-expressing U251 and PC3 cells (*Figure 4A, B and Figure 4—figure supplement 1A–E*). DNA synthesis still took place in some of the monstrous nuclei 5 days after switching on SHC016 expression. This was demonstrated by fluorescence microscopy analysis of incorporation of BrdU, added at the end of U251 culture, into DNA *(Figure 4—figure supplement 1F)* and was in agreement with cytometric analysis of BrdU/viability staining, which showed a significant fraction of BrdU-incorporating cells (~32%) among SHC016-expressing U251 (Figure 2B). However, BrdU was not always evenly distributed within DNA, indicating unsynchronized DNA replication in some SHC016-expressing U251 cells *(Figure 4—figure supplement 1F).*

The images of interphase nuclei of SHC016-expressing cells additionally revealed irregular patches of heterochromatin (which is also a trait of necrosis and senescence), numerous micronuclei, and in some cases supernumerary centrosomes (*Figure 4C and Figure 4—figure supplement 2*). During metaphase the formation of multipolar spindle occurred much more often in SHC016-expressing U251 than in SHC002-expressing U251 or in either cell line cultured without doxycycline. Infrequently, the defects in chromosome alignment were also observed in U251 with induced expression of SHC016 (*Figure 4D, E*).

The analysis of the expression of genes coding for proteins involved in the cell cycle execution and control including these responsible for correct spindle formation showed that the expression of SHC016 in U251 did not markedly affect the levels of most transcripts, at least up to 48 h of doxycycline induction (*Figure 4—figure supplement 3*). Interestingly, similarly to other cell lines tested, the levels of both *SFN* and *GADD45A* transcripts decreased in response to SHC016 expression in U251. However, since the changes in the expression of these genes do not apply to absolutely all tested cells and are not always rapid, we rejected the idea that either of them may play a role of a primary mediator of SHC016-induced effects.

In search for this common first trigger we performed the screening RNAseq analysis of RNA isolated from two cell lines, A549 and U251 transduced with Tet-on-SHC016, incubated for 24 h with- or without doxycycline. Three genes, whose levels were substantially diminished in both cell lines after inducing SHC016 expression caught our attention: *ARPP19, VPS4B,* and *SNRPD3.* The activity of ARPP19, cAMP-regulated phosphoprotein 19, which is an inhibitor of PP2A phosphatase, is required for maintaining high levels of cyclin B1-CDK1 complex during mitosis (Gharbi-Ayachi et al., 2010). VPS4B, vacuolar protein sorting B, is a part of ESCRT III complex involved in cytokinesis (Morita et al., 2010) and SNRPD3, small nuclear ribonucleoprotein Sm D3, is a component of U1, U2, U4, and U5 spliceosomal ribonucleoprotein (snRNP) complexes (Will & Luhrmann, 2011).

To verify the importance of RNAseq results we analyzed, via RT-qPCR, the influence of SHC016 expression on *ARPP19*, *VPS4A, VPS4B*, and *SNRPD3* in all studied human cell lines and in murine MC38CEA. Although RNAseq analysis has shown decrease in *VPS4A* expression only in A549 cells we included this gene because VPS4A is essential for proper spindle formation and cytokinesis (Morita et al., 2010), the processes which are seriously disturbed in U251 cells. The levels of *ARPP19*, *VPS4B*, and *SNRPD3* transcripts were decreased in all human cell lines in response to SHC016-, but not to SHC002 expression, and the most spectacular drop was observed for *SNRPD3 (Figure 5A* and *Figure 5—figure supplement 1*). The levels of *VPS4A* were diminished in response to SHC016 in all human cell lines but MCF7. In murine MC38CEA cells expressing SHC016, the levels of *Arpp19* and *Vps4b* were not affected; the expression of *Vps4a* was moderately- and this of *Snrpd3* – strongly suppressed *(Figure 5—figure supplement 1).*

**Figure 5.**
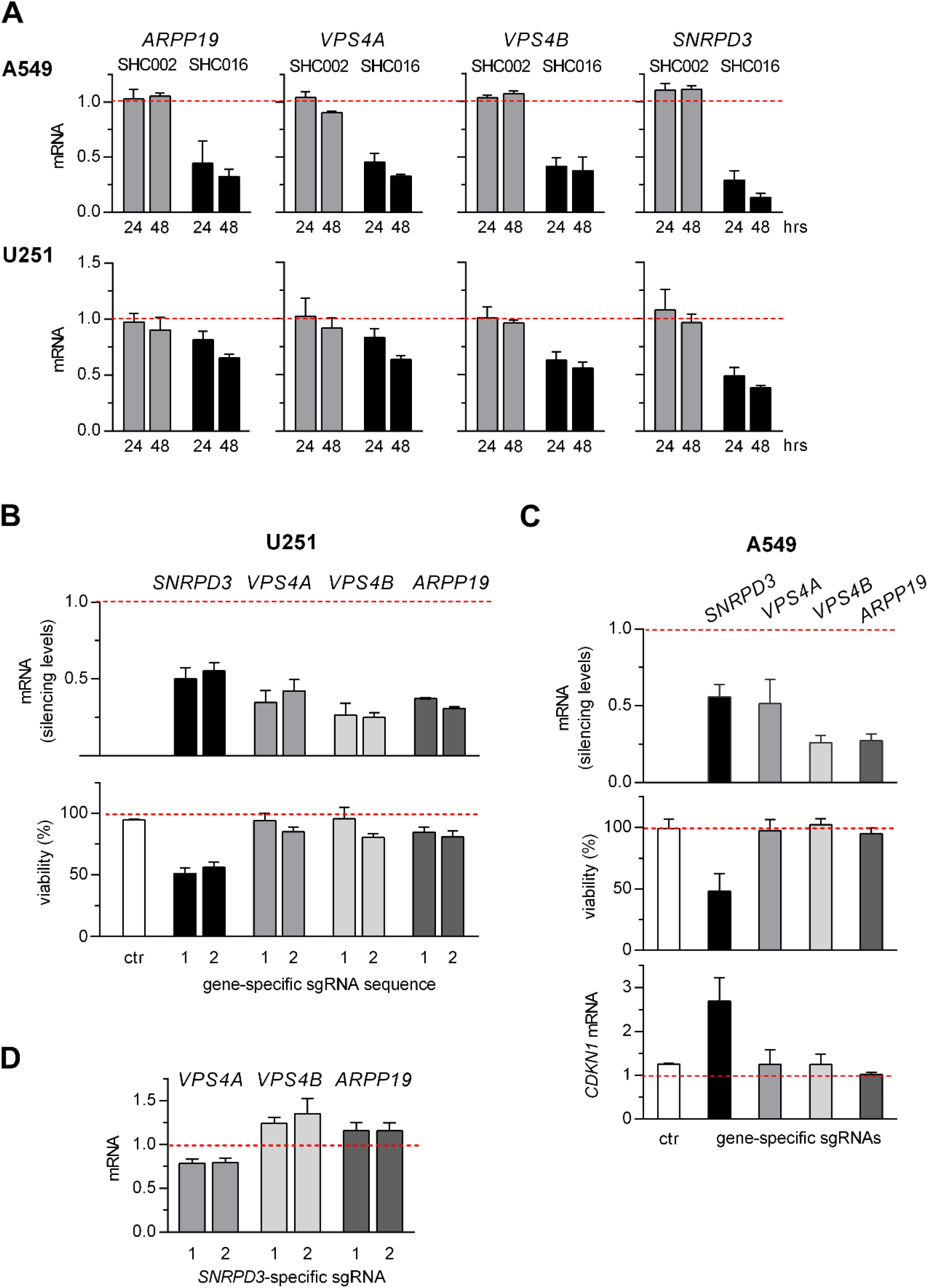
Deleterious effects of SHC016 are caused by diminished *SNRPD3* transcript levels. (A) RT-qPCR analysis of transcripts identified by RNAseq as potentially affected by SHC016 performed 24- and 48 h after inducing expression of SHC002 or SHC016 shRNAs in A549 and U251 cells. Mean values ± SD of relative mRNA levels from three independent experiments are shown. The data concerning other cell lines are available in *Figure 5—figure supplement 1*. (B) The analysis of extents of *SNRPD3, VPS4A, VPS4B,* and *ARPP19* silencing and viability of U251 cells in which expression of particular genes was diminished by CRISPRi using two different sgRNAs for each gene (designated as 1 and 2; the sequences are presented in *Figure 5— supplementary table 1*). Detailed data concerning the results of gene silencing are presented in *Figure 5—figure supplement 2*. (C) The analysis of extents of *SNRPD3*, *VPS4A*, *VPS4B*, and *ARPP19* silencing in A549 cells in correlation with cell viability (assessed by MTT) and *CDKN1* expression (assessed by RT-qPCR). Two sgRNAs (1/2) were used simultaneously in CRISPRi-mediated gene silencing. (B, C) MTT was performed on the sixth day after inducing dCas-KRAB-MeCP2 expression. Ctr – control cells transfected (U251) or electroporated (A549) with an empty CRISPRi plasmid (pSBtet-Pur-dCas9-KRAB-MeCP2-hU6-SapI), which does not contain a sequence coding for any gene-targeting sgRNA. (D) RT-qPCR analysis of potential influence of *SNRPD3*-silencing on *VPS4A*, *VPS4B*, and *ARPP19* expression in U251 cells. (B-D) RNA was isolated 72 h after inducing dCas-KRAB-MeCP2 expression. (A-D) The relative levels of the transcripts in uninduced cells were taken as 1. MV ± SD from three independent experiments are shown.

To answer the question whether the decrease in the expression of any of these genes may mimic the effects of SHC016 we applied a doxycycline-inducible CRISPRi system in A549 and U251 cells using two different sgRNAs for the silencing of each gene transcription. Because CRISPRi inhibits *de novo* RNA synthesis and, unlike shRNA, does not affect existing RNAs, therefore we analyzed the effects induced by CRISPRi later than those of shRNAs; the levels of particular mRNAs were measured 72 h- and the effects on cell viability was examined 6 days after inducing dCas-KRAB-MeCP2 expression.

In U251 cells, the levels of the transcripts were decreased only moderately when doxycycline was used at a concentration of 100 ng/ml to induce dCas9-KRAB-MeCP2 expression *(Figure 5—figure supplement 2*). At higher doxycycline concentration (1 μg/ml), the levels of the transcripts were reduced to around 50% for *SNRPD3,* 40% for *VPS4A,* 25% for *VPS4B,* and 35% for *ARPP19* of their initial levels *(Figure 5B* and *Figure 5—figure supplement 2).* Remarkably, lowering the expression of *SNRPD3* in U251 cells only by half resulted already in a strong decrease in cell viability (*Figure 5B*) associated with hallmarks of mitotic catastrophe, observed previously in SHC016-expressing U251 cells *(Figure 5—figure supplement 3*). The silencing of either *VPS4A* or *VPS4B,* or *ARPP19* had little effect on the viability of U251 cells *(Figure 5B)* and nuclear morphology (data not shown).

Unfortunately, A549 cells were resistant to transfection and the attempts to efficiently silence the expression of all genes of interest via cell transfection with CRISPRi plasmids bearing one sgRNA sequence failed *(Figure 5—figure supplement 4A).* Simultaneous transfection of A549 cells with two CRISPRi plasmids coding for different gene-specific sgRNAs improved the outcome but still the levels of *SNRPD3* and *VPS4A* transcripts were diminished only by ~30% *(Figure 5—figure supplement 4A)*. When, in this scheme, transfection was replaced by electroporation the levels of silencing reached ~50% for *SNRPD3* and *VPS4A,* and ~75% for *VPS4B* and *ARPP19 (Figure 5— figure supplement 4A*). The decrease in *SNRPD3* mRNA level, but not of any other studied transcript, resulted in a substantial decline in A549 viability and increase in *CDKN1* expression (*Figure 5C*). Interestingly, even small reductions in *SNRPD3* mRNA levels observed after transfection of A549 with a single or double CRISPRi plasmids were accompanied by diminished viability of the cells (*Figure 5—figure supplement 4B*).

We concluded, that the reduced levels of *SNRPD3* in response to SHC016 expression account for the processes that lead to death or at least to permanent growth arrest of various cells transduced with the MISSION^®^ non-targeting SHC016 vector. Because SNRPD3 is a subunit of Sm protein complex involved in splicing, we speculated that its insufficiency may result in the defective splicing of *VPS4A*, *VPS4B*, or *ARPP19* transcripts and decrease in the levels of these mRNAs. However, in U251 cells, CRISPRi-diminished expression of *SNRPD3* was not accompanied by diminished but rather slightly increased levels of *VPS4B* and *ARPP19* transcripts. In contrast, the levels of *VPS4A* was moderately reduced upon *SNRPD3* silencing *(Figure 5D).*

How is it possible that the scientists around the world widely use the MISSION^®^ system without noticing the effects we here describe? Usually, the commercially available vectors with noninducible expression of shRNAs are used and therefore the initial massive death of cells is being attributed to puromycin effect on non-transduced cells. We believe that with subsequent cell divisions only cells with low expression of SHC016 survive. Even in the case of the cells transduced with the vector coding for doxycycline-inducible SHC016, discontinuation of the use of selective antibiotic, puromycin, being in line with generally accepted protocols, resulted in less deleterious effects of SHC016 as compared to the cells, in which SHC016 was induced with doxycycline in the presence of puromycin. The viability of MCF7 and HeLa cells, evaluated with MTT test was further reduced by almost 20 percent points, when SHC016-expressing cells were cultured for 5 days in the presence of puromycin *(Figure 6A).* Also, addition of puromycin to the culture medium of HeLa cells caused the increase in population of dead cells from 5% to 40% in SHC016-but not in SHC002-expressing cells, as measured by BrdU/viability test *(Figure 6B* and *Figure 6—figure supplement 1*).

**Figure 6.**
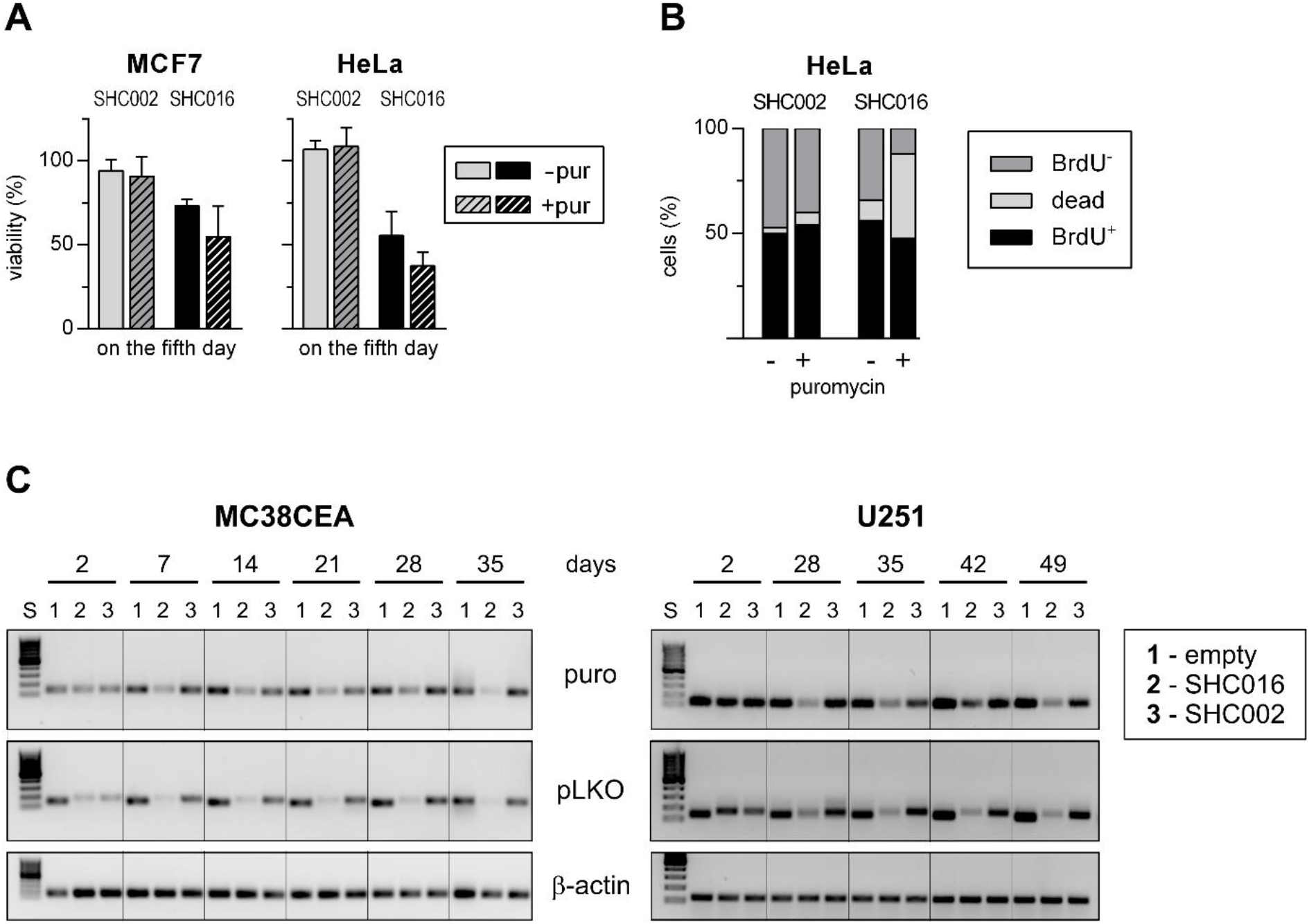
SHC016 expression cassette is eliminated from the cell cultures. (A) Comparison of the viability of MCF7 and HeLa cells 5 days after induction of either SHC002 or SHC016 shRNA expression in the absence or presence of selection antibiotic, puromycin (1 μg/ml), which was added for the last 72 h of culture. Absorbance values of the cells without induction of shRNA expression were taken as 100%. Data are shown as MV ± SD from 3 independent experiments. (B) Comparison of DNA synthesis and cell viability assessed via BrdU/eFluor 520 viability staining of HeLa cells expressing for 5 days either SHC002 or SHC016 in the absence or presence of selection antibiotic, puromycin. Dead – all eFluor 520^+^ cells, BrdU^+^ – eFluor 520^−^/BrdU^+^ cells, BrdU^−^ – eFluor 520^−^/ BrdU^−^ cells. Different representations of the data containing error bars can be found in *Figure 6—-figure supplement 1.* (C) PCR analysis of the abundance of the elements of pLKO vectors in the long term cultures of MC38CEA and U251 cells. The cells were transduced with the empty vector (SHC001), or SHC002, or SHC016. DNA was isolated after indicated times and the sequence from puromycin-resistance gene (puro) and the sequence comprising shRNA insertion site (pLKO) were amplified. Inverted images of DNA electrophoresis gels are shown. S – size standard, GeneRuler 100 bp DNA Ladder, Thermo Scientific. The amplified fragments are as follow: puro – 123 bp, pLKO empty – 136 bp and pLKO coding for either non-targeting RNA – 170 bp.

To verify the hypothesis that only the cells with low expression of SHC016 may remain in a longterm culture we transduced two cell lines, MC38CEA and U251, with uninducible, original versions of MISSION^®^ control vectors: empty vector (SHC001) and two vectors with constitutive expression of the non-targeting shRNA sequences, SHC002 and SHC016. We isolated DNA from the cultured cells two days after transduction and then at one-week intervals and analyzed the levels of the transgenes by PCR using two pairs of primers, one amplifying the sequence comprising PGK promoter and puromycin resistance gene and the second flanking the shRNA cloning site. Puromycin was added 48 h after transduction; the cells were kept under its selective pressure for the next seven days. In the case of U251 cells, very high mortality of the cells transduced with pLKO-SHC016 did not allow to perform the analysis earlier than in a month after transduction. In either cell line the levels of SHC002 and SHC016 cassettes were similar in two days after transduction. However, in contrast to the empty vector and to pLKO-SHC002, the level of the transgene containing SHC016 decreased dramatically during the cell propagation *(Figure 6C).*

## Discussion

The original enthusiasm accompanying the increasing use of RNAi techniques for exploring gene functions has declined in recent years with the emergence of publications pointing to their drawbacks and limitations, such as inefficient silencing of target gene expression, which may lead to false-negative results and—much more dangerous for the further application of the analyses— false-positive off-target effects (Housden & Perrimon, 2016). Already “classic” publication by Jackson et al. (2003) demonstrated that different siRNA targeting the same transcript show unique global gene expression profiles with only few genes regulated in common (Jackson et al., 2003). Commercial availability of shRNA libraries and individual shRNA sets which silence almost every known gene have reduced the verifiability of experimental results from various laboratories. Falsepositive results due to unintended silencing of off-target genes were positively verified by results obtained in other laboratories that used the same silencing sequences. This may have far-reaching, disastrous consequences because RNA interference-based studies are often the cornerstones for development of novel therapies. The excellent study by Shelzer’s group provided crushing evidence for misdirected drug development based on erroneous RNAi data leading to failures in clinical trials (Lin et al., 2019). They tested 5 proteins (HDAC6, MAPK14, PAK4, PBK, and PIM1), selected, based mostly on RNAi data, as the so-called cancer dependencies, i.e. genes that encode proteins required for the survival and/or proliferation of cancer cells, and for which small molecule drugs that specifically block their effects were already in clinical trials or preclinical studies. Using CRISPR-based techniques the authors proved that neither knockout (via CRISPRko) nor reduction of the expression level (via CRISPRi) of those genes affected the viability of 32 various cancer cell lines. These genes were apparently wrongly selected probably due to RNAi off-target effects. Interestingly, an OTS964 inhibitor selected as an inhibitor of one of these targets, namely PBK, did exhibit potent antimitotic activity but by inhibiting different kinase, CDK11B. Thus, a completely different group of cancer patients is the target group for treatment with this compound than it would appear from RNAi-based studies (Lin et al., 2019; Lin, Giuliano, Sayles, & Sheltzer, 2017).

These examples and also other critical analyses evidence that off-target effects may be wrongly attributed to the analyzed genes. Our work points to another, as yet overlooked, possible cause of discrepancies between various genotype-to-phenotype data and frequent failures of verification of RNAi results by competitive or complementary techniques. The RNAi results may be also misinterpreted due to the off-target effects related to the so-called non-targeting shRNA control, which by definition should not significantly affect the expression of any gene. What are the consequences of silencing activity of a non-targeting shRNA for interpretation of RNAi data?

The first to come to mind is that the inhibition of a gene expression caused by a control shRNA, which plays a role of a reference, would be erroneously interpreted as the stimulation of the expression of that gene by experimental shRNAs.

However, in the case of SHC016, the control shRNA applied in our work and probably widely used in other studies employing MISSION^®^ shRNAs, the outcome is more complex because SHC016 reduces the expression of a gene essential for cell survival, namely *SNRPD3*. During posttransduction antibiotic treatment, most likely selected are the cells with highly reduced expression of this toxic shRNA. Therefore, the levels of non-targeting and targeting shRNAs are incomparable and the cells expressing non-targeting shRNA cease to function as a proper control.

As shRNAs utilize miRNA processing machinery and compete with pre-miRNA for Dicer and Ago proteins (Kim, Kim, & Kim, 2016; Ma, Zhang, & Wu, 2014), disparate loads of shRNAs in control and experimental cells may distinctly influence endogenous miRNA functions, which will manifest by differences in gene expression profiles unrelated to specific shRNA effects. Moreover, during antibiotic treatment, in our case puromycin, selected are the cells which survive in the presence of antibiotic despite reduced occurrence of puromycin *N*-acetyltransferase cassette associated with the low levels of SHC016 coding sequence. This functional phenotype can be attributed to the cells overexpressing multidrug resistance gene(s) coding for ABC transporters able to expel puromycin from the cells (Theile, Staffen, & Weiss, 2010). Apart from the differences in the expression levels of endogenous gene(s) rendering puromycin resistance also unmatched intracellular levels of transgenic puromycin *N*-acetyltransferase and/or puromycin itself may distinctly affect cell transcriptomes (Guo, Lee, Byrnes, & Miller, 2017).

It might seem that applying more restrictive rules for designing of control- and gene-specific shRNAs could limit off-target effects, but the problem is quite complex. Today, shRNA off-targets are still hardly predictable, because of alternative selection of siRNA strands, lower complementarity requirements than initially assumed, and variations in shRNA-derived siRNAs. Argonaute protein (Ago) determines which strand of a miRNA duplex will serve as a silencing guide strand and supposedly favors the one whose 5’ sequence (the first four nucleotides) comprises less thermodynamically stable end of the duplex and whose first nucleotide is preferentially U (>A>>C and G). However, only ~25% of human miRNA are chosen accordingly to both conditions and as much as ~25% totally eludes these rules (Medley, Panzade, & Zinovyeva, 2020). Likewise, the strand of shRNA-derived siRNA designed to be a passenger strand can also be loaded into Ago (Sheng et al., 2020), thus doubling the number of potential off-targets. Although sh- and siRNAs are designed to have an array of 19-20 nucleotides complementary to sequences within specific targets, the complementarity of a seed region comprising nucleotides 2-7/8 of a guide strand and a hexamer or heptamer in a 3’UTR of mRNA may be sufficient for off-target activity (Birmingham et al., 2006). However, other patterns of complementarity between effectively silencing miRNAs and target mRNA sequences refuted even this requirement (Akbari Moqadam, Pieters, & den Boer, 2013; Brennecke, Stark, Russell, & Cohen, 2005). Additionally, the existence of isomiRs, the variants of miRNA derived from the same pre-miRNA, which slightly differ in length and/or identity of terminal nucleotide(s) (Gebert & MacRae, 2019) suggests that also shRNAs may be subject to inaccurate processing. In accordance with this assumption one nucleotide shifts was observed in processing of pTIP- and pLKO-based shRNAs (Putzbach et al., 2017). More pronounced deviations from the designed shRNA-derived silencing sequences were revealed by Bhinder et al. who demonstrated non-canonical Dicer-independent processing of shRNA encoded by a TRC vector, so possessing the same hairpin scaffold as SHC016. They showed that *CTTN-* targeting shRNA is able to silence also six other nonrelated genes due to generation of various siRNA comprising nucleotides of 5’ or 3’ arm as well as those of a loop and termination signal (Bhinder et al., 2014). Imprecise processing of TRC shRNAs was also reported in other studies (Gu et al., 2012; Watanabe, Cuellar, & Haley, 2016).

Human *SNRPD3* has two translatable transcript variants, which differ only in 5’UTR, both with two alternative polyadenylation sites. There is one known mouse *Snrpd3* transcript variant with short 3’UTR. It shows 92% and 61% sequence identity of CDS and 3’UTR, respectively, with corresponding regions of both human variants according to the results of Needleman-Wunsch alignment (Altschul et al., 1997). Because both human and mouse transcript were silenced by SHC016 we assumed that homologous sequences within these mRNAs should interact with SHC016-derived siRNA. Taking into account the undeniable silencing of *SNRPD3* and still undefined rules governing the silencing process, we did not focus on identifying transcripts’ and shRNA regions responsible for this effect.

SNRPD3 (small nuclear ribonucleoprotein D3) belongs to Sm core proteins of the spliceosome and plays pivotal roles in all steps of splicing. Given that more than 95% of human protein-coding transcripts are subject to splicing, it is reasonable to expect that insufficiency of any of spliceosome core proteins would have global detrimental effects on cells and organisms. Surprisingly, the mutations of various spliceosome core proteins do not lead to generalized splicing defects but are rather linked with different events of alternative splicing (Vazquez-Arango & O’Reilly, 2018). Growing body of evidence indicates, that apart from trans-acting regulatory splice factors, also spliceosome core proteins are responsible for the pattern of alternative splicing. For example Papasaikas et al., while analyzing the splicing pattern of transcripts coding for proteins involved in apoptosis or cell cycle, revealed that the silencing of spliceosomal core proteins, including SNRPD3, showed a stronger tendency to promote exon skipping than the noncore ones (Papasaikas, Tejedor, Vigevani, & Valcarcel, 2015). The differences in splicing landscape in response to the deficiency of particular core splice factors may reflect the differences in the strength of intron splice sites in combination with the expression profile of splicing regulators.

Nevertheless, we did not observe global reduction in transcript levels in response to *SNRPD3* silencing and thus we suspected that the gradual decrease of this protein level primarily affects the expression of a few proteins crucial for cell cycle progression.

*SNRPD3* was selected in genomic screenings as one of the genes, whose depletion cause lethal defects in key steps of mitosis including metaphase chromosome alignment (Hofmann, Husedzinovic, & Gruss, 2010) and sister chromatid cohesion (Sundaramoorthy, Vazquez-Novelle, Lekomtsev, Howell, & Petronczki, 2014). However, not much is known about possible transcripts coding for proteins crucial for cell cycle progression, whose splicing might depend on SNRPD3 levels and whose defective splicing might explain the effects observed by us upon silencing of *SNRPD3* expression either unintentionally via SHC016 or purposely using CRISPRi.

One candidate is a centromere protein E (CENP-E) transcript. CENP-E, a plus-end-directed kinesin-7 motor protein, is involved in chromosome congression and microtubule-kinetochore conjugation as well as in activation of spindle assembly checkpoint. Its deficiency may lead to mitotic arrest followed by cell death. It has been shown that enhancer rudimentary homolog (ERH), which is required for *CENPE* splicing, interacts with SNRPD3 (Weng et al., 2012). Silencing of SNRPD3, similarly to silencing of ERH in DLD-1 and HCT116 colorectal cancer cell lines led to diminished expression of *CENPE.* It resulted, as confirmed for ERH-silenced cells, from nonsense mediated decay of defectively spliced *CENPE* transcript (Weng et al., 2012). We observed that *CENPE* expression was strongly diminished in A549 (~80%) but only slightly (~15%) in U251 cells 48 h after inducing SHC016 expression (*Discussion—supplement 1*). As *CENPE* is one of DREAM targets, the reduction of its mRNA level in A549 cells, the timing and extent of which resembles those of other DREAM-repressed genes, most probably results simply from p53 activation.

The lack of substantial decrease of *CENPE* mRNA in U251 cells 48 h after inducing SHC016 expression suggests that *CENPE* is not responsible for the early manifestations of mitotic catastrophe triggered by *SNRPD3* silencing in U251 cells. However, because the half-life of SNRPD3 (protein) was estimated to be > 27 h (Boisvert et al., 2012), it is possible that a gradual reduction in *CENPE* mRNA levels would occur over time. Indeed, in early time points (48 h after induction), centrosome amplification seems to be the culprit for mitotic catastrophe observed in U251 cells, because defects in chromosome congression accounted only for a small part of all mitotic defects. However, centrosome staining on day 5 after induction of shRNA expression revealed that only a part of cells that had undergone mitotic catastrophe had supernumerary centrosomes *(Figure 4C* and *Figure 4—figure supplement 2*), pointing out yet another mechanism that drives aberrant mitosis and, in consequence, mitotic catastrophe. In fact, after a longer period of SHC016 expression, defects in chromosome congression were prevalent (data not shown) and the changes in gene expression profile at later time points (up to 5 days) in *SNRPD3*-silenced, p53-null cells would be required to fully explain the molecular basis of the observed effects.

*SNRPD3* was also selected in genome-wide screening for genes, whose silencing strongly influence the viability of p53-wild type cancer cells (Siebring-van Olst et al., 2017). The authors demonstrated that the silencing of several splicing factors, including SNRPD3, induced prominent activation of p53 in A549 non-small cell lung cancer cells, which is in agreement with our data showing activation of p53-p21 axis in A549 cells following SHC016 expression. They further show that cytotoxicity mediated by *SNRPD3* silencing did not depend on p53 activity, which is again in line with our observations of deleterious effects of SHC016- or CRISPRi-mediated silencing of *SNRPD3* both in cells proficient or deficient in active p53. It has also been demonstrated that silencing of *SNRPD3* promoted *MDM4* exon 6 skipping leading to the expression of a shorter and more labile transcript variant (Siebring-van Olst et al., 2017), which in turn could affect the level of MDM4 and p53-dependent and p53-independent functions of the protein (Haupt, Mejia-Hernandez, Vijayakumaran, Keam, & Haupt, 2019). For example MDM4 has recently been shown to facilitate unperturbed DNA replication via suppressing the formation of RNA-DNA hybrids (Wohlberedt et al., 2020). We observed substantially diminished *Mdm4* mRNA levels in SHC016-expressing cells but only in those of murine origin (Figure 1). Neither in A549 nor U251 cells the levels of *MDM4* transcript were affected 48 h after inducing SHC016 expression (data not shown). Additionally, given only a slight decrease in BrdU incorporation and the type of aberrations in p53-mutant U251 and PC3 cells indicating that mitosis and not DNA replication are affected we believe that at least in these cells the altered splicing of transcript(s) other than *MDM4* plays a crucial role in mediating effects of SHC016 expression and *SNRPD3* silencing. Recently, another protein, namely structural proteasome subunit β3, whose transcript *PSMB3* was defectively spliced in SNRPD3-deficient A549 cells was suggested as a culprit of cell lethality (Blijlevens et al., 2020). Silencing of SNRPD3 or other Sm protein coding genes resulted in a prevalence of the expression of a long variant of *PSMB3* associated with significantly reduced proteasome activity (Blijlevens et al., 2020). This may have far-reaching consequences as the cell cycle relies on the timely proteasome-driven degradation of factors involved in subsequent stages of the cycle. Proteolysis is particularly critical at the metaphase-to-anaphase transition as destruction of securin and cyclin B are necessary for sister-chromatid separation and mitotic exit (Morgan, 2007). Interestingly, diminished proteasomal activity might explain the appearance of supernumerary centrosomes with subsequent multipolar spindle formation in U251 cells. It has been shown, for example, that excess accumulation of MPS1 (Monopolar Spindle 1) spared from proteasomal degradation drive centriole overduplication (Kasbek et al., 2007). Also pharmacologic inhibition of the proteasome mimics the effect of overexpression of certain proteins involved in centriole duplication, which is manifested by formation of multiple daughter centrioles from a single mother centriole (Brownlee & Rogers, 2013). Thus our results fits the scenario that at least some effects of *SNRPD3* silencing results from diminished activity of proteasome, although verification of the hypothesis requires further studies.

Taking into account the key role of SNRPD3 in splicing, most probably many transcripts including those coding for proteins important for cell survival and proliferation are affected by incorrect alternative splicing. In our studies expression of SHC016 shRNA resulted not only in silencing of *SNRPD3* but also in reduced levels of other mRNAs that may influence cell cycle progression but are not DREAM complex targets. Based on qPCR analysis we rejected the hypothesis that diminished levels of *VPS4B* and *ARPP19* as well as those of *Sfn/SFN* and *Gadd45A/GADD45A* observed in SHC016 expressing cells are the aftermath of aberrant splicing associated with *SNRPD3* silencing *(Figure 5D* and data not shown, respectively). Thus either SHC016 affects their stability directly or influences their levels indirectly but independently of SNRPD3 status. It remains to be verified whether slight changes in the transcript levels of multi-intronic *VPS4A* and *VPS4B* in U251 cells with CRISPRi-diminished expression of *SNRPD3* is associated with alterations in splicing events.

In summary: SHC016 expression lowers the level of at least several transcripts encoding proteins essential for successful cell cycle execution. Of these, only *SNRPD3* silencing has a detrimental effect on all tested cells.

SNRPD3 has been indicated as a potential target in cancer therapy (Blijlevens et al., 2020; Siebring-van Olst et al., 2017). Similarly, down-regulation of other Sm proteins, functional partners for SNRPD3, was also more detrimental for tumor cells than for normal ones (Blijlevens, van der Meulen-Muileman, de Menezes, Smit, & van Beusechem, 2019; Putzbach et al., 2017). Our research confirms strong dependence of cancer cell survival on undisturbed SNRPD3 activity. In our hands, however, MEF, non-tumor murine cell line, is also sensitive to SHC016, which affects *SNRPD3* level. MEFs are immortalized, embryonal, rapidly proliferating cells, thus they do not represent normal, primary cells and their sensitivity does not discredit core spliceosome proteins as cancer therapeutic targets. However, in-depth studies should be undertaken to determine whether differences in sensitivity of cancer cells and normal proliferating cells, e.g. intestine epithelium or immune cells to reduced levels of Sm proteins justify considering them as potential therapeutic targets.

The example of SHC016, which targets SNRPD3, a gene essential for cell survival, highlights that control shRNAs may elicit strong off-target effects that invalidate reliable analysis of RNAi data. Despite continued efforts to improve shRNA scaffolds (Sheng et al., 2020), the chances for elimination or significant reduction of RNAi off-target activity while preserving on-target efficiency are low. Processing of shRNA and its interactions with binding partners are influenced by many poorly understood variables, but most importantly siRNA mimics a natural miRNA mechanism, which by definition does not show high specificity – one miRNA usually silences from several to several dozen transcripts (Liu & Wang, 2019).

In recent years CRISPR/Cas9-based methods (CRISPRko and CRISPRi) evolved as an alternative for RNAi (Housden & Perrimon, 2016; Schuster et al., 2019). Similarly to shRNA libraries, also libraries of sgRNAs suitable for either CRISPR/Cas9-based high-throughput analyses are available (Schuster et al., 2019). CRISPR/Cas9-based methods are not flawless. First of all, the genetic background of analyzed cells is modified if Cas endonuclease or Cas-fusion protein are delivered to the cells by introducing a gene coding for either one. Secondly, the efficiency of CRISPRko may be affected by cell ploidy and locus accessibility, while CRISPRi requires annotated transcription start sites (TSS) and is not a method of choice for genes transcribed from more than one TSS. However, CRISPR/Cas9-based methods outcompete RNAi in terms of minimizing off-target effects (Doench et al., 2016; Evers et al., 2016; Naeem, Majeed, Hoque, & Ahmad, 2020). Development of high-fidelity Cas enzymes greatly reduced off-target activity of Cas9 (Kleinstiver et al., 2016; Slaymaker et al., 2016) reported at early stage of CRISPRko technology. The situation is even better for CRISPRi due to the very mechanism of action. The efficient inhibition of transcription requires long-term and high-affinity interaction between 17-20 nucleotide guide sequence and target DNA and therefore mismatch tolerance is low and the number of off-targets – negligible (Gilbert et al., 2014; Peddle, Fry, McClements, & MacLaren, 2020).

In light of our research, we believe that the use of SHC016 as a control non-targeting sequence should be discontinued. Moreover, all data of RNAi analyses based on this control should be reevaluated, reinterpreted, and verified using other methods including e.g. CRISPR/Cas9-based techniques. Our work also supports the notion that all RNAi data that were not verified by complementary methods should be treated with caution and scientific skepticism (Morgan, 2007). Rapid development of methods utilizing the extraordinary properties of sgRNA and Cas endonuclease complexes has a potential to dethrone the RNAi technique, making it an auxiliary method in genetic loss-of-function research.

## Materials and methods

### Cell lines culture

The following cell lines were used: MC38CEA (murine colon adenocarcinoma (Corbett, Griswold, Roberts, Peckham, & Schabel, 1975) expressing human carcinoembryonic antigen (Bereta et al., 2007)), MEF (mouse embryonic fibroblasts, a gift from Prof. Paul Saftig, Christian-Albrechts University Kiel, Germany), U251 MG (human glioblastoma, verified by Eurofins Medigenomics, Ebersberg, Germany), A549 (human lung carcinoma, ATCC CCL-185), PC3 (human prostate adenocarcinoma, ATCC CRL-1435), MCF7 (human breast adenocarcinoma, ATCC HTB-22), and HeLa (human cervical adenocarcinoma, ATCC CCL-2). HeLa cells were grown in MEM (Lonza, Basel, Switzerland) and all other cell lines in DMEM (Corning Inc, Corning, NY), supplemented with 10% heat-inactivated fetal bovine serum (FBS), tetracycline negative (Capricorn Scientific, Ebsdorfergrund, Germany) at standard conditions. The cell cultures were routinely tested by PCR for mycoplasma contamination using mycoplasma rDNA-specific primers.

### Lentiviral Vectors

MISSION^®^ pLKO.1-puro Empty Vector Control Plasmid DNA, which does not contain any shRNA insert (SHC001), MISSION^®^ pLKO.1-puro Non-Mammalian shRNA Control Plasmid DNA (SHC002), containing shRNA insert, which, according to description, does not target any known mammalian genes, MISSION^®^ pLKO.1-puro Non-Target shRNA Control Plasmid DNA (SHC016), containing shRNA insert, which, according to description, does not target any genes from any species. To produce doxycycline-inducible shRNA expression vectors, oligonucleotides coding for non-targeting shRNAs, identical to those in MISSION SHC002 and SHC016, CCGGCAACAAGATGAAGAGCACCAACTCGAGTTGGTGCTCTTCATCTTGTTGTTTTT and CCGGGCGCGATAGCGCTAATAATTTCTCGAGAAATTATTAGCGCTATCGCGCTTTTT, respectively, were annealed and cloned into *Age*I/*Eco*RI-linearized tet-pLKO-puro (Addgene plasmid #21915, a gift from Dmitri Wiederschain (Wiederschain et al., 2009)). A resulting vectors are referred to as Tet-on-SHC002 and Tet-on-SHC016.

### Production and titration of lentiviral vectors

Two days prior to transfection, 293T cells were plated on 10 cm plates. The cells were transfected with plasmids: 1.3 pmol psPAX2, 0.72 pmol pMD2.G (Addgene plasmids #12260 and #12259; both were gifts from Didier Trono) and 1.64 pmol respective pLKO plasmid using PEI MAX 40K (Polysciences, Inc, Warrington, PA) at a ratio of DNA to PEI 1:3. Medium was changed 4 h after transfection. Cell culture supernatants containing pseudoviral particles were collected 48 h later, filtered through 0.45 μm PES filters and concentrated by overnight centrifugation at 8500 g, 4°C. Pellets containing pseudoviral particles were resuspended in equal volumes of serum-free DMEM. Initially, vectors were titrated using QuickTiter Lentivirus Titer Kit (Lentivirus-Associated HIV p24; Cell Biolabs, San Diego, CA); the viral titers for SHC002 and SHC016 were similar, therefore in subsequent experiments equal volumes of concentrated viruses were used.

### Cell transduction

The cells were grown in a 12-well plate. Optimal volume of viruses was determined experimentally by transducing each cell line with several dilutions of a concentrated viral stock. Aliquots of 2 μl of stocks, which resulted in 20-60% of puromycin-resistant cells, were eventually used for transduction of all cell types via spinoculation (30 min, 1150×g, room temperature, RT) in the presence of polybrene (8 μg/ml). After 48 h the transduced cells were selected for 7 days with puromycin (Bioshop Canada Inc, Burlington, Canada) at a concentration of 1 μg/ml for human cell lines, 5 μg/ml for MC38CEA and 8 μg/ml for MEF.

### PCR analysis of stability of transgene integration

MC38CEA and U251 MG (further referred to as U251) cells transduced with lentiviral vectors, SHC001, SHC002, and SHC016 were cultured for several weeks. DNA was isolated from the cells at weekly intervals. The cells were lysed in guanidinium thiocyanate solution and DNA was isolated by phenol-chloroform extraction.

Equal amounts of DNA samples (100 ng) were subjected to PCR using Taq Master Mix (Vazyme Biotech, Nanjing, China) to amplify puromycin resistance gene, puromycin N-acetyl-transferase, and a pLKO fragment comprising shRNA coding sequence as well as actin coding sequence *(ACTB/Actb)* as a control. The primers are listed in Manuscript supplement—Table I. PCR program included 30 cycles of 30 s at 94°C, 30 s at 60°C (human and mouse actin) or 52°C (puro and pLKO), 30 s at 72°C. PCR products were visualized on a 1% agarose gel.

### Stimulation of shRNA expression

The cells: MC38CEA, MEF, U251, A549, PC3, MCF7, and HeLa transduced with Tet-on-SHC002 or Tet-on-SHC016 were seeded at a density appropriate for each cell line (between 500 – 1500 cells/well in a 96-well plates or 10,000 – 20,000 cells/well in a 12-well plates). The cells were cultured for 5 or 6 days and the cells of each line were divided into several experimental groups. Starting one day after plating of the cells, doxycycline aliquots were added every 24 h to one experimental group to a final concentration of 100 ng/ml (unless stated otherwise). One group (negative control) was left untreated. In some experiments etoposide (2 μM) was added to one group for the last 48 h of culture as a positive control of apoptosis induction.

### MTT assay

The viability of the cells was determined by MTT assay. The cells in 96-well plates were incubated in 100 μl of serum-free medium containing thiazolyl blue tetrazolium bromide (MTT, Sigma-Aldrich, St. Louis, MO) at a concentration of 0.5 mg/ml for 1–3 hours. Formazan crystals were solubilized in 200 μl of acidified isopropanol. The absorbance was measured at 570 nm using a microplate reader (VersaMax, Molecular Devices, San Jose, CA). The absorbance of control cells (cultured without doxycycline) was taken as 100% viability.

### Colony formation assay

The cells were seeded at 100 cells per well in a 6-well plates. On the following day doxycycline was added at 100 ng/ml. The cells were cultured for 7–10 days and fresh doxycycline was added every other day. At fifth day puromycin was added to select the cells that retained the expression cassette. The colonies were stained with crystal violet dissolved in methanol, destained with tap water and photographed with Fusion FX imaging platform (Vilber Lourmat, Collégien, France).

### Annexin V assay

The cells were analyzed by an annexin V/propidium iodide double-staining method (Vermes, Haanen, Steffens-Nakken, & Reutelingsperger, 1995). The cells with inducible SHC002 or SHC016 expression were grown in 12-well plates and shRNA expression was induced with doxycycline as described above. The cells were trypsinized, washed with PBS and then with Annexin V Binding Buffer (ABB: 10 mM HEPES pH 7.4, 140 mM NaCl, 2.5 mM CaCl2), and incubated in ABB containing Annexin V conjugated with APC (1% v/v; Exbio, Vestec, Czechia) and PI (100 μg/ml) for 20 min in dark. Next, the cells were washed twice with ABB, resuspended in ABB and analyzed on FACS Calibur (BD Biosciences, San Jose, CA). The percentages of apoptotic cells (annexin V-positive, PI-negative) and late apoptotic and necrotic cells (PI-positive, annexin V-positive and negative) were determined using FlowJo v10.0.7 (FlowJo LLC, Ashland, OR).

### Measurement of caspase activity

MC38CEA and MEF cells with inducible SHC002 or SHC016 expression were seeded in 96-well plates and shRNA expression was induced with doxycycline as described above. Caspase activity was determined using Caspase-Glo^®^ 3/7 Assay Systems (Promega, Madison, WI) according to the manufacturer’s specifications. The luminescence was measured using a microplate reader Synergy H1 Hybrid Reader (BioTek, Winooski, VT). The luminescence values of control cells, non-treated with doxycycline, were taken as 100%.

### Cell cycle analysis

The cells with inducible SHC002 or SHC016 expression were plated in 12-well plates at a density of 10,000 cell/well and shRNA expression was induced with doxycycline as described above. The cells were trypsinized, washed with PBS, fixed with 70% ethanol, and incubated at 4°C for at least 4 h. Equal amounts of cells (1×10^5^), were permeabilized with 0.1% Triton X-100 in PBS for 15 min, washed with PBS, and incubated in 200 μl of ribonuclease A (10 μg/ml, A&A Biotechnology, Gdynia, Poland) for 15 min at 37°C. Then 200 μl aliquots of propidium iodide (PI) dissolved in PBS containing 0.1% Triton X-100 were added to the cells to a final concentration of 250 μg/ml. The cell cycle analysis of 10,000 cells/sample was performed by flow cytometry using FACS Calibur (BD Biosciences) and FlowJo cell cycle Watson (Pragmatic) model (FlowJo v10.0.7 software, FlowJo LLC). Both debris and doublets were removed from the analysis.

### Nuclei imaging

The cells were washed twice with PBS, fixed with ice-cold methanol for 5 min, washed thoroughly with PBS, incubated with DAPI (1 μg/ml) or Hoechst 33342 (1 μg/ml) for 15 min, and washed 4 times with PBS. The nuclei were imaged using a Leica DM6 B microscope equipped with Leica DMC5400 camera or DM IL LED fluorescence microscope equipped with Leica DFC450 C (Leica Microsystems, Wetzlar, Germany) camera and the images were processed using ImageJ software version 1.53 (National Institutes of Health) (Rueden et al., 2017; Schindelin et al., 2012).

### BrdU labelling

The cells were seeded at a density of 20,000 cells/well (all but A549) or 60,000 cells/well (A549) in a 6-well plate. On the following day, doxycycline was added to a final concentration of 100 ng/ml. The cells (all but A549) were cultured for 5 days or 3 days (A549). Fresh doxycycline was added every other day. BrdU (Sigma-Aldrich), at a final concentration of 20 μM, was added to the medium for the last six hours, then the cells were collected, washed twice with PBS and stained with Fixable Viability Dye eFluor 520 (Thermo Scientific, Waltham, MA) in PBS for 30 min at 4°C with occasional mixing. To stop the reaction, the cells were washed once in 1% BSA/PBS and once in PBS. Then, the cells were fixed dropwise with ice cold 70% ethanol and incubated overnight at 4°C. On the next day, the fixed cells were subjected to DNA denaturation in 2 M HCl containing 0.5% Triton X-100 for 30 min at RT; HCl was neutralized with 0.1 M sodium tetraborate, pH 8.5. After extensive washes in PBS, the cells were incubated in blocking solution (1% BSA in PBST) for 20 min and then with Alexa Fluor 647-conjugated anti-BrdU antibody at 1:50 (clone MoBU-1, Thermo Scientific) for at least 3 h at RT. Next, the cells were centrifuged, suspended in PBS and analyzed on FACS Calibur (BD Biosciences). The percentages of BrdU positive, BrdU negative and dead cells (Fixable Viability Dye eFluor 520-positive) were determined using FlowJo v10.6.1 (FlowJo LLC). In some experiments U251 cells were also seeded on #1.5 glass coverslips and stained as describe above with minor modification (viability dye was omitted; staining was performed with Alexa Fluor 647 or Alexa Fluor 488-conjugated anti-BrdU antibodies). The samples were analyzed via fluorescence microscopy using Leica DM6 B microscope equipped with 63×1.3 NA oil objective and Leica DMC5400 camera (Leica Microsystems).

### Senescence-associated β-galactosidase staining

The cells were seeded 5 days prior to staining at 5000 cells/well in 12-well plates and shRNA expression was induced with doxycycline as described above. SA-β-gal staining was performed according to a standard protocol (Debacq-Chainiaux, Erusalimsky, Campisi, & Toussaint, 2009). Briefly, the cells were washed twice with PBS, and fixed for 5 min with 2% formaldehyde and 0.2% glutaraldehyde in PBS, then washed in PBS and incubated overnight at 37°C in staining solution containing 40 mM citric acid/sodium phosphate buffer, pH 6.0, 1 mg/ml X-Gal (Bioshop Canada Inc), 5 mM potassium ferricyanide, 5 mM potassium ferrocyanide, 150 mM NaCl and 2 mM MgCl_2_. On the following day, the cells were washed with PBS, dehydrated in methanol, counterstained with hematoxylin and washed extensively with water. The cells were imaged using a DM IL LED microscope equipped with Leica DFC450 C camera. The percentage of SA-β-gal-positive cells was calculated. About 300 cells were analyzed per experiment in three independent experiments.

### RNA isolation, reverse transcription and RT-qPCR

Total RNA was isolated by the modified phenol/chloroform extraction method using Fenozol reagent (A&A Biotechnology), then treated with TURBO DNase (Thermo Scientific) followed by column purification with Clean Up RNA Concentrator (A&A Biotechnology). Equal amounts of RNA (1 μg) were reverse-transcribed with a mixture of oligo(dT)15 and random hexamer primers using M-MLV polymerase (Promega). RT-qPCR gene expression quantifications were performed using AceQ qPCR SYBR Green Mix (Vazyme Biotech) on Eco Real-Time PCR System (Illumina, San Diego, CA). RNA expression was normalized to a geometric mean of two reference genes: *Eef2* and *Polr2b* for mouse cell lines and *EEF2* and *TBP* for human cell lines. All sequences of pimers are listed in Manuscript supplement—Table I.

### Preparation of samples for RNA sequencing

Cells were seeded on 10 cm plates two days before doxycycline addition. After 24 h of doxycycline treatment, cells were lysed in RNA Extracol (EurX, Gdańsk, Poland). Total RNA was isolated using Direct-zol RNA Mini (Zymo Research, Irvine, CA); the procedure included on-column DNase I digestion. Poly(A)^+^ fractions were isolated using Dynabead mRNA DIRECT Micro Purification Kit (Thermo Scientific) according to manufacturer recommendations with minor modifications such as three (instead of two) rounds of RNA binding, washing and elution and additional wash in detergent-free buffer B before final elution step. Libraries preparation and sequencing on Ion Torrent platform was performed at Genomics Centre at Malopolska Centre of Biotechnology (Kraków, Poland).

### Construction of tet-inducible CRISPRi vector

dCas9 repressor was PCR-amplified with primers containing *Sfi*I sites from dCas9-KRAB-MeCP2 (a gift from Alejandro Chavez & George Church; Addgene plasmid #110821) (Yeo et al., 2018) digested with *SfiI* and cloned into *Sfi*I-linearized pSBtet-Pur plasmid (a gift from Eric Kowarz; Addgene plasmid #60507) (Kowarz, Loscher, & Marschalek, 2015).

In order to clone U6 promoter-sgRNA scaffold cassette containing *Sap*I restriction sites instead of *Bbs*I sites, oligonucleotides containing *Sap*I sites were annealed and cloned into *Bbs*I-digested pX330-U6-Chimeric_BB-CBh-hSpCas9 (a gift from Feng Zhang; Addgene plasmid #42230) (Cong et al., 2013). U6-sgRNA scaffold was then PCR-amplified with primers containing *KpnI* sites and cloned into *Kpn*I-digested pSBtet-Pur-dCas9-KRAB-MeCP2. The resulting vector was termed pSBtet-Pur-dCas9-KRAB-MeCP2-hU6-SapI.

sgRNA sequences targeting human *SNRPD3, VPS4A, VPS4B,* and *ARPP19* genes were designed using GPP sgRNA Designer (Broad Institute). Oligonucleotides containing pSBtet-Pur-dCas9-KRAB-MeCP2-hU6-SapI compatible ends were annealed and assembled with the vector in one restriction-ligation reaction with *Sap*I and T4 DNA ligase.

U251 and A549 were seeded onto 12-well plates one day prior to transfection. The cells were transfected with 475 ng of a respective CRISPRi plasmid together with 25 ng of Sleeping Beauty transposase-encoding vector pCMV(CAT)T7-SB100 (a gift from Zsuzsanna Izsvak; Addgene plasmid #34879) (Mates et al., 2009) using jetOPTIMUS (U251 cells; Polyplus-transfection, Illkirch, France) or TransIT-LT1 (A549 cells; Mirus Bio, Madison, WI) according to the protocols provided by the manufacturers. Alternatively, A549 cells were electroporated with 4.9 μg of CRISPRi plasmid and 100 ng of pCMV(CAT)T7-SB100 in electroporation buffer (100 mM sodium phosphate, 10 mM MgCl_2_, 5 mM KCl, 20 mM HEPES, 50 mM sodium succinate; pH 7.2) using Gene Pulser II (Bio-Rad Laboratories, Inc., Hercules, CA) under following conditions: two million cells in 400 μl electroporation buffer in 0.4 cm gap cuvette, pulse voltage and capacitance: 300 V and 1000 μF.

### Immunofluorescence

The cells were plated on #1.5 glass coverslips in 12-well plates. On the following day, doxycycline was added to a final concentration of 100 ng/ml and 48 h later, the cells were washed with PBS, fixed with ice-cold methanol for 5 min, then washed three times with PBS, permeabilized with 0.5% Triton X-100 in PBS for 15 min at RT, blocked with 2% BSA and 5% normal goat serum in PBST (blocking solution) for 1 h, and incubated with mouse anti-γ-tubulin antibody (clone C-11, 2 μg/ml, Santa Cruz Biotechnology, Dallas, TX) diluted in blocking solution overnight at 4°C.

After washing with PBST, the coverslips were incubated with Fab fragment of goat anti-mouse IgG (H+L) secondary antibody conjugated to Alexa Fluor-594 (Jackson Immunoresearch, West Grove, PA) for 1 h at RT. After extensive washing with PBST, the coverslips were subjected to mild fixation with 0.4% methanol-free formaldehyde in PBS for 10 min, washed with PBS and incubated with CoraLite-488-conjugated mouse anti-β-tubulin antibody (clone 1D4A4, 10 μg/ml; Proteintech, Rosemont, IL) in blocking solution overnight at 4°C. After washing with PBST, the coverslips were stained with Hoechst 33342 and mounted onto microscopic slides with ProLong Glass (Thermo Scientific). When the cells were stained for γ-tubulin only, the post-staining fixation step was omitted. The images were acquired with Leica DM6 B microscope equipped with Leica DMC5400 camera using 63×1.3 NA oil objective. Images shown are maximum intensity projections of z-stack planes.

## Additional information

Data analysis was performed using Microsoft Excel (Excel 2016) or GraphPad Prism v. 5 and all graphs were created using GraphPad Prism v. 6 (GraphPad Software).

## Acknowledgements

This work was funded by the National Science Centre, Poland (NCN, OPUS11 funding scheme, project number UMO-2016/21/B/NZ3/00355 to J.B.). We thank dr. Krystyna Stalińska and Ms. Olga Dracz for help in some experiments and Dr. Mateusz Wawro for critically reading the manuscript.

## Supplementary data

### Figure 1.

Expression of SHC016 in MC38CEA results in apoptosis and cell cycle arrest at the G2/M phase.

**Figure supplement 1A.**
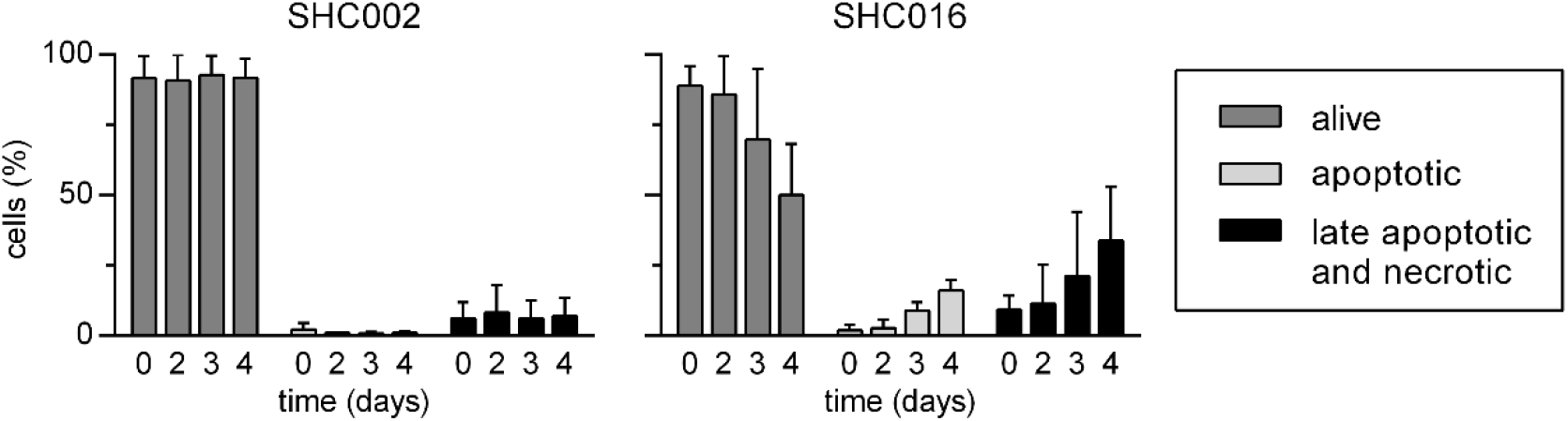
Expanded version of Figure 1B of the main text. Analysis of apoptotic death of MC38CEA cells expressing SHC002 or SHC016 assessed by annexin V/PI staining and flow cytometry analysis; alive cells – annexin V^−^PI^−^; apoptotic – annexin V^+^/PI^−^; late apoptotic and necrotic – annexin V^+^/PI+ and annexin V^−^/PI^+^ (the latter usually did not exceed 3% of the cells expressing either shRNA). MV ± SD from three independent experiments are shown.

**Figure supplement 1B.**
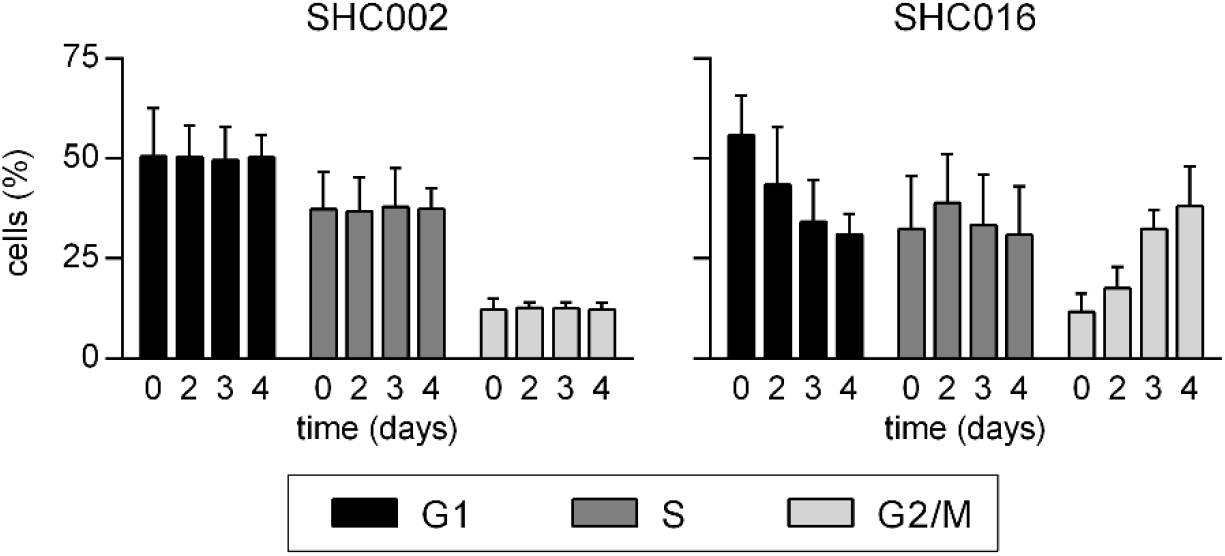
Expanded version of Figure 1D of the main text. Flow cytometry analysis of the cell cycle of MC38CEA cells expressing SHC002 or SHC016. MV ± SD from three independent experiments are shown.

**Figure supplement 2.**
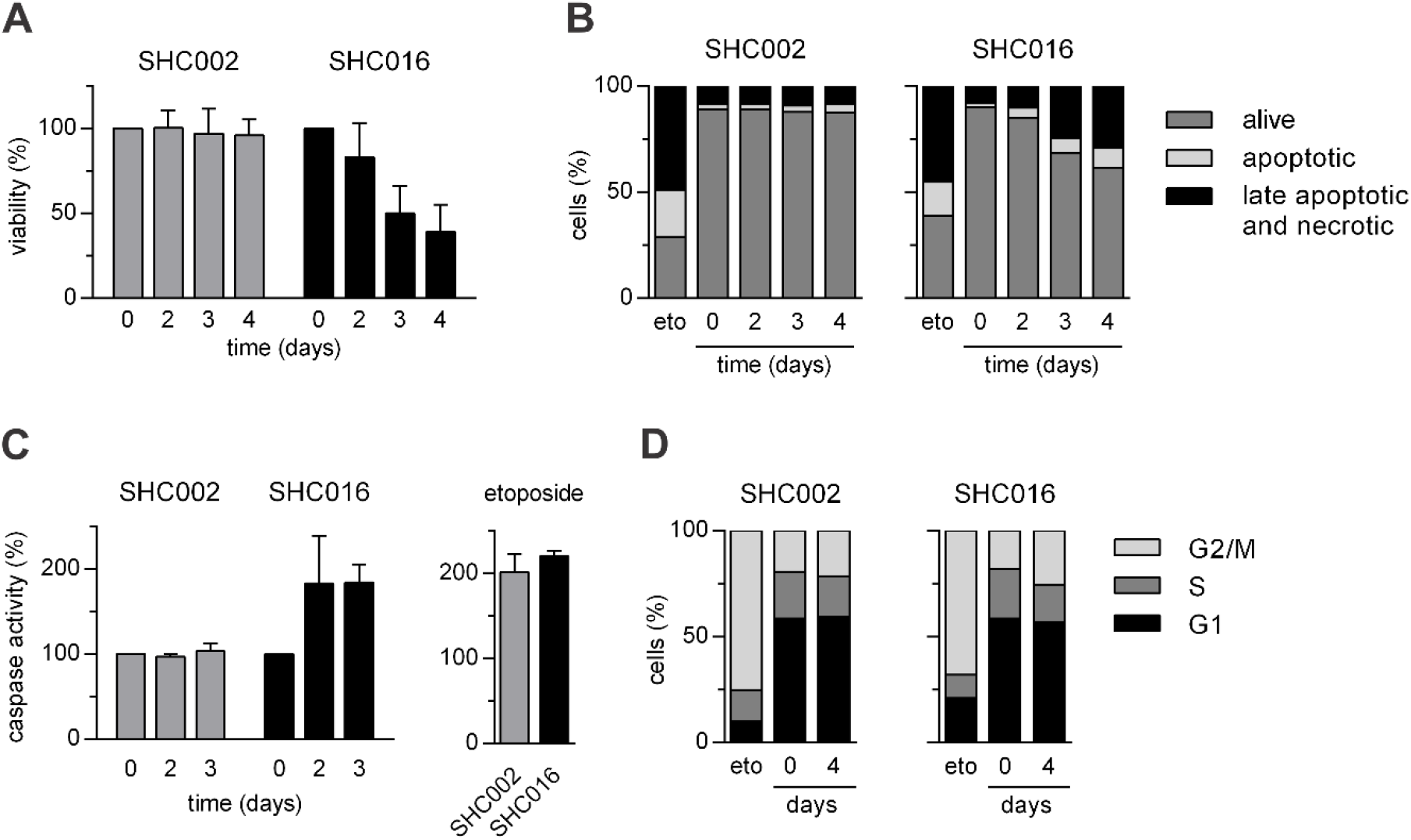
Expression of SHC016 in MEF cell results in apoptosis. The expression of non-coding shRNA SHC002 or SHC016 was induced by doxycycline (100 ng/ml) for the last 4, 3, or 2 days of the 5-day culture. (A) Cell viability assessed by MTT assay. Absorbance values of the cells with uninduced transgene expression were taken as 100%. (B) Death via apoptosis assessed by annexin V/PI staining and flow cytometry analysis; alive cells – annexin V^−^PI^−^; apoptotic – annexin V^+^/PI^−^; late apoptotic and necrotic – annexin V^+^/PI^+^ and annexin V^−^/PI^+^. (C) Activity of caspases 3 and 7 measured via chemiluminescence assay. Chemiluminescence values of the cells with uninduced transgene expression were taken as 100%. (D) Flow cytometry analysis of the cell cycle. (B, C, D) The cells treated for the last 2 days of experiment with etoposide (2 μM) were used as a positive control. Data are shown as MV ± SD from 3 independent experiments.

**Figure supplement 3.**
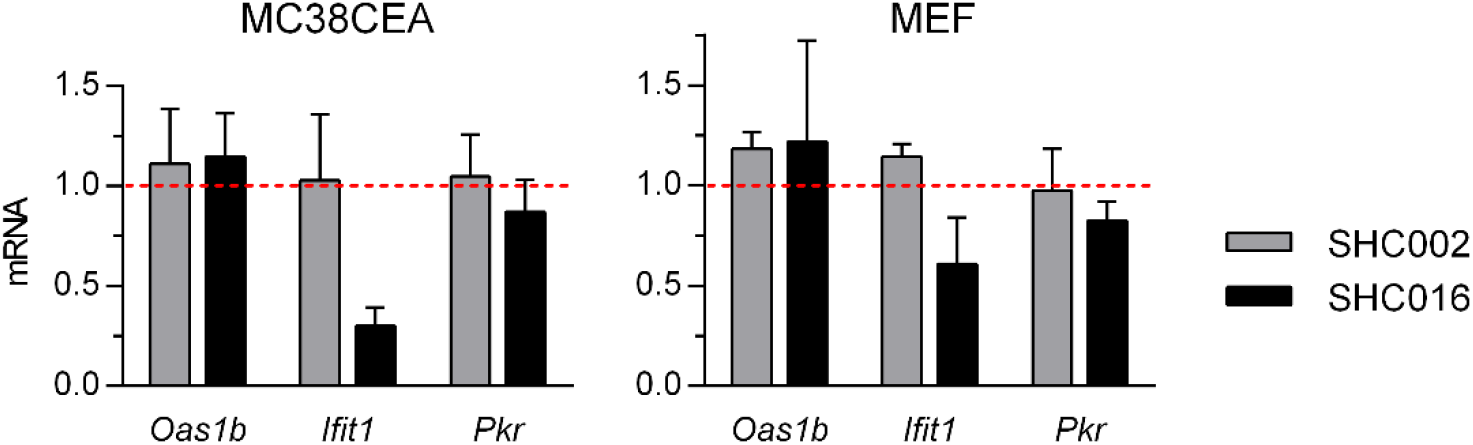
Expression of non-coding RNAs, SHC002 or SHC016, did not induce interferon response in murine cells. RT-qPCR analysis demonstrated that the levels of three IFN-inducible transcripts were not augmented 48 h after inducing SHC002 or SHC016 expression. The relative levels of the transcripts in uninduced cells were taken as 1. Data are shown as MV ± SD from 3 independent experiments.

### Figure 2

**Figure supplement 1.**
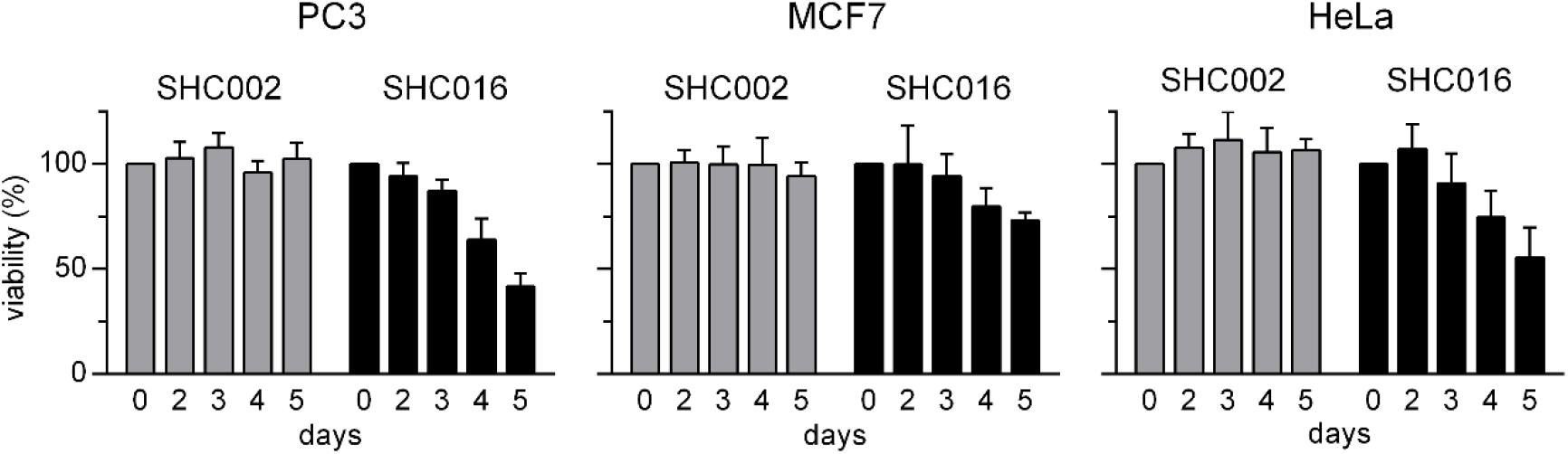
Expanded version of data presented in Figure 2A of the main text. Cell viability assessed by MTT assay. The expression of non-coding shRNA SHC002 or SHC016 was induced by doxycycline (dox, 100 ng/ml) for the last 5, 4, 3, or 2 days of the 6-day culture. Absorbance values of the cells with uninduced transgene expression were taken as 100%. MV ± SD of three independent experiments are shown.

**Figure supplement 2.**
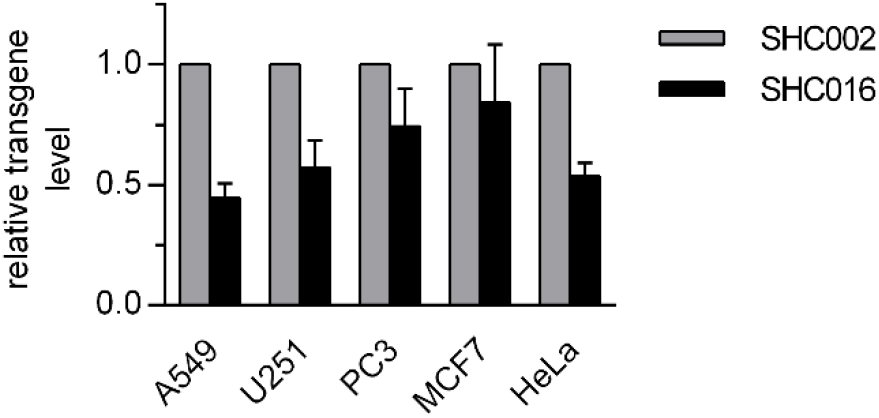
qPCR analysis of the levels of SHC002 and SHC016 transgenes incorporated into genomes of the studied cell lines. The cells were transduced with Tet-on-SHC002 or Tet-on-SHC016 vector. DNA was isolated using Blood/Cell DNA Mini Kit (Syngen Biotech) after primary puromycin selection. Equal amounts of DNA (15 ng) were subjected to qPCR using AceQ Universal SYBR Green Master Mix (Vazyme Biotech) on Eco Real-Time PCR System (Illumina). PCR program included DNA denaturation at 95°C for 5 min, followed by 40 cycles of 10 s at 95°C and 30 s at 60°C. All samples were analyzed in duplicates. H1-pLKO cassette level was normalized to human genomic locus LINC00657 level utilizing the ΔΔCT method with correction for primers efficiency. Primers’ sequences are available in *Manuscript supplement—Table I.* MV ± SD of three (or four in the case of PC3 cells) independent experiments are shown. The relative level of SHC002 in each cell line was taken as 1.

**Figure supplement 3.**
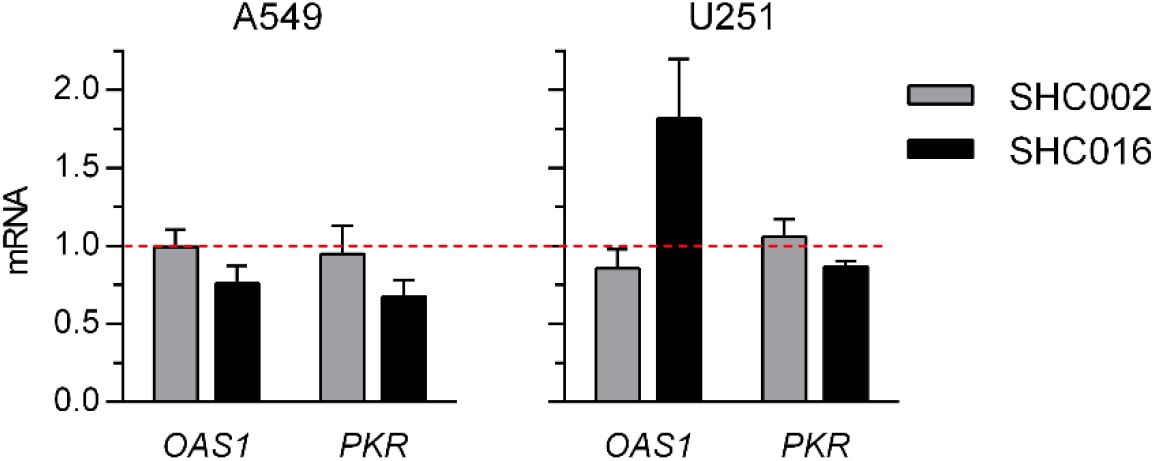
Expression of non-coding shRNAs, SHC002 or SHC016, do not induce interferon response in human A549 and U251 cell lines. RT-qPCR analysis performed 48 h after inducing SHC002 or SHC016 expression demonstrated that the levels of two IFN-inducible transcripts were not augmented to the levels characteristic for interferon response, which is between 10 and 10^4^ times in the case of *OAS1^1^*. The relative levels of the transcripts in uninduced cells were taken as 1. MV ± SD of three independent experiments are shown.

**Figure supplement 4.**
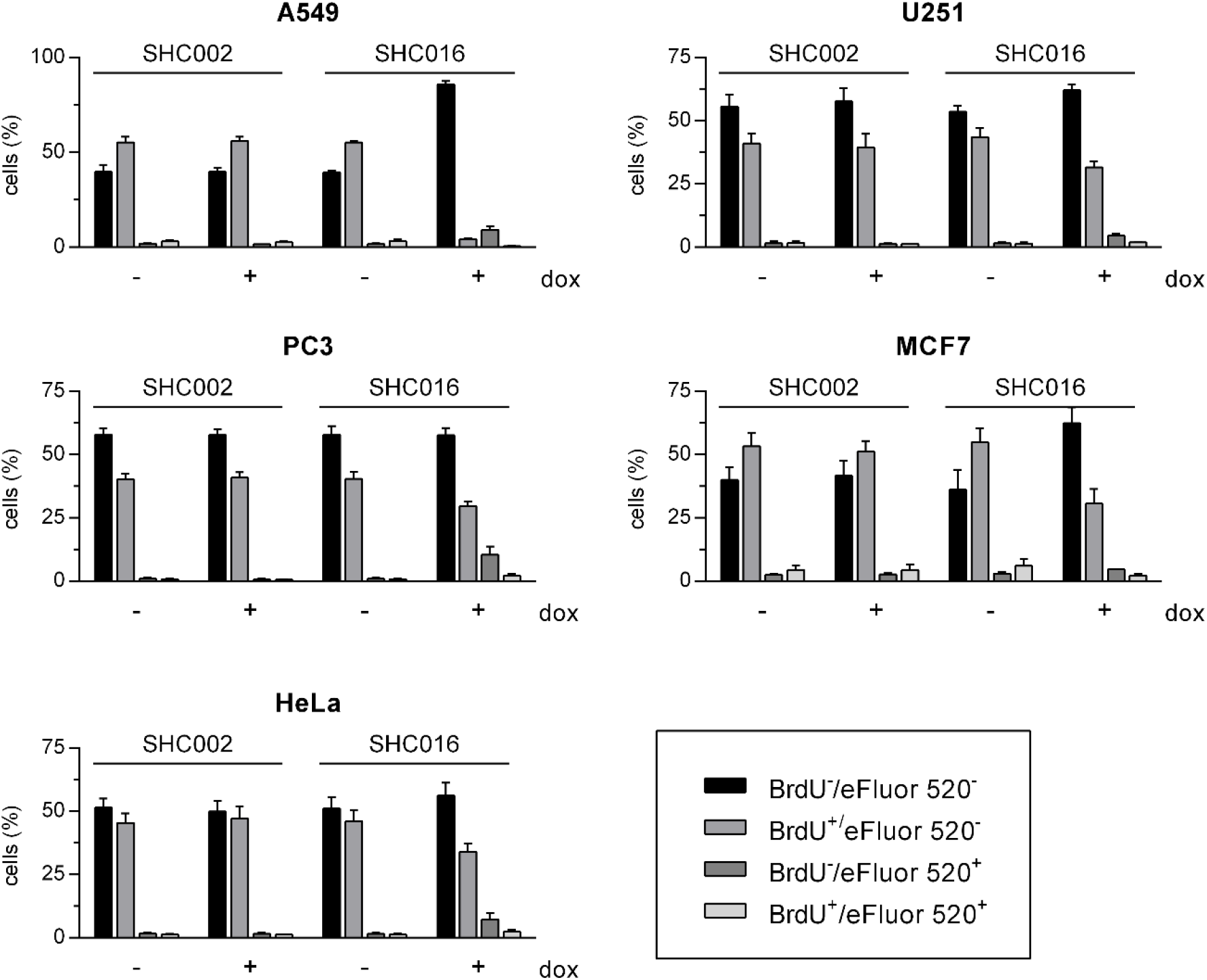
Expanded version of data presented in Figure 2B of the main text. DNA synthesis and cell viability assessed via double BrdU/eFluor 520 viability staining performed three (A549) or five days (U251, PC3, MCF7, HeLa) after inducing SHC002- or SHC016 expression. The difference of the time schedule was dictated by shorter time needed to manifest cytostatic/cytotoxic effects of SHC016 in A549 and substantially shorter doubling time of A549 than of other cell lines (*Figure 2—figure* supplementary table I). BrdU was added to the medium for the last 6 h of culture. MV ± SD of three independent experiments are shown.

**Supplementary Table I.** Estimation of population doubling time of the studied cell lines

| Cell line* | | Doubling time<br>(MV ± SD)** |
| --- | --- | --- |
| A549 | SHC002 | 13.55 ± 1.39 |
| | SHC016 | 13.45 ± 1.48 |
| U251 | SHC002 | 20.10 ± 7.53 |
| | SHC016 | 19.27 ± 6.33 |
| PC3 | SHC002 | 19.60 ± 3.67 |
| | SHC016 | 19.40 ± 3.39 |
| MCF7 | SHC002 | 27.93 ± 3.91 |
| | SHC016 | 23.70 ± 3.30 |
| HeLa | SHC002 | 21.76 ± 1.49 |
| | SHC016 | 20.97 ± 1.65 |

\*Doubling time was estimated in **uninduced** cell cultures. The cells were seeded in 24-well plates ( $5 \times 10^3$ per well) in DMEM supplemented with 10% FCS. The cells were trypsinised and counted every 24 h for 5 days using a Bürker chamber or an automated cell counter (TC20, BioRad).
\*\*Population doubling time ( $T_d$ ) was calculated in the logarithmic growth phase by the formula:
$T_d = (t_2 - t_1) \frac{\ln 2}{\ln(q_2/q_1)}$ ; $t_1$ and $t_2$ are initial and final times and $q_1$ and $q_2$ are initial and final cell numbers, respectively. MV ± SD from three (A549, U251, and MCF7) or two (PC3, HeLa) independent experiments are presented.

#### Figure 3

**Figure supplement 1.**
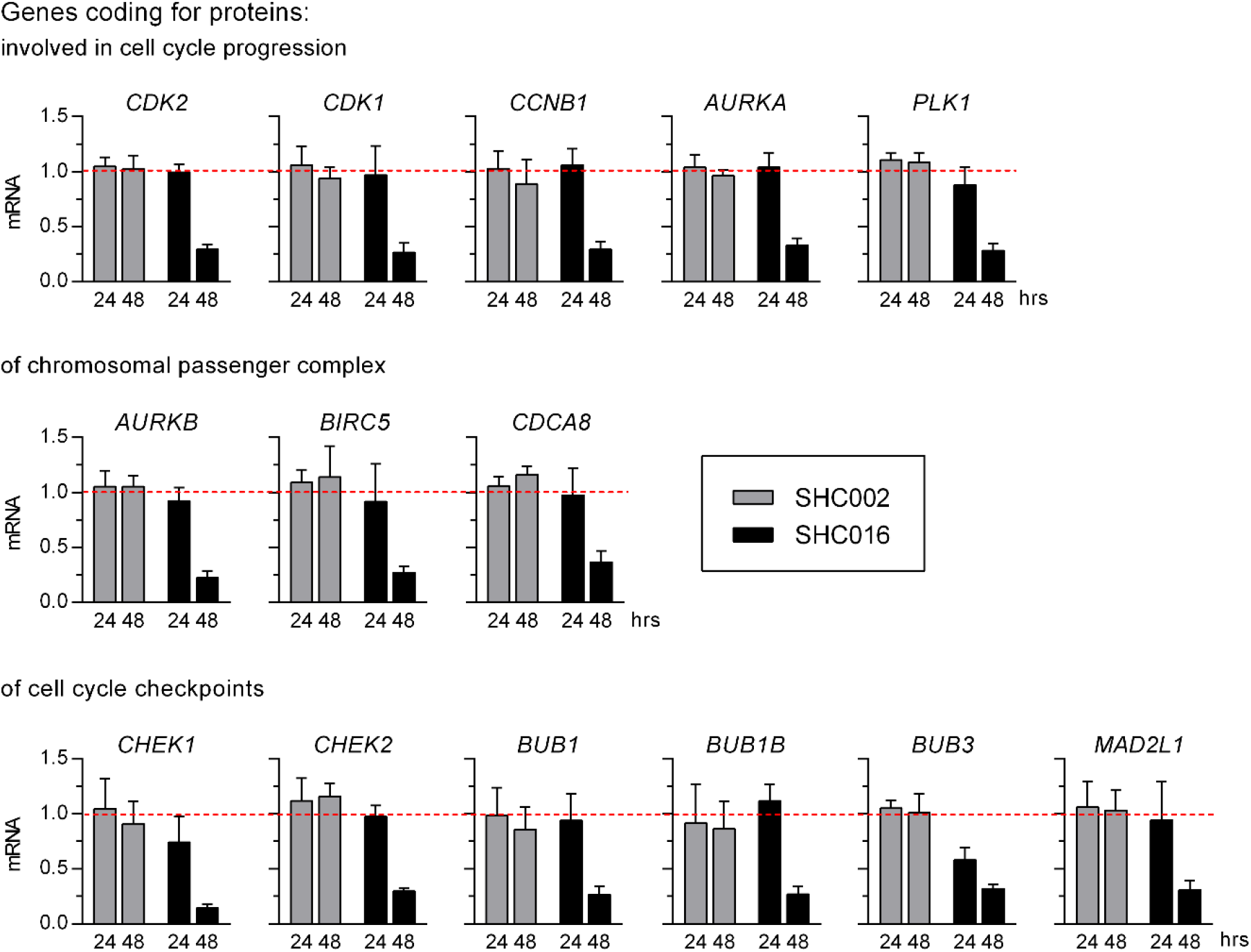
A549 cells were subjected to RT-qPCR analysis of mRNA levels of genes involved in cell cycle progression and control. mRNA was isolated 24 or 48 h after inducing expression of SHC002 or SHC016 shRNAs. The relative levels of the transcripts in uninduced cells were taken as 1. Primers’ sequences are available in *Manuscript supplement—Table I.* MV ± SD from 3 independent experiments are shown.

**Figure supplement 12.**
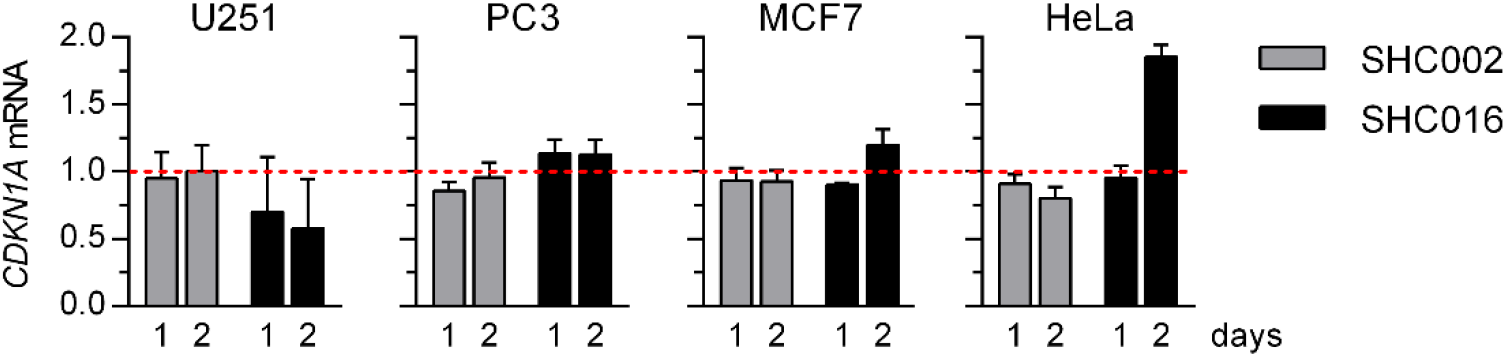
RT-qPCR analysis of *CDKN1A* mRNA levels in the studied cell lines 24 or 48 h (1 or 2 days) after inducing expression of SHC002 or SHC016. The relative levels of the transcript in uninduced cells were taken as 1. MV ± SD from 3 independent experiments are shown.

#### Figure 4

**Figure supplement 1A.**
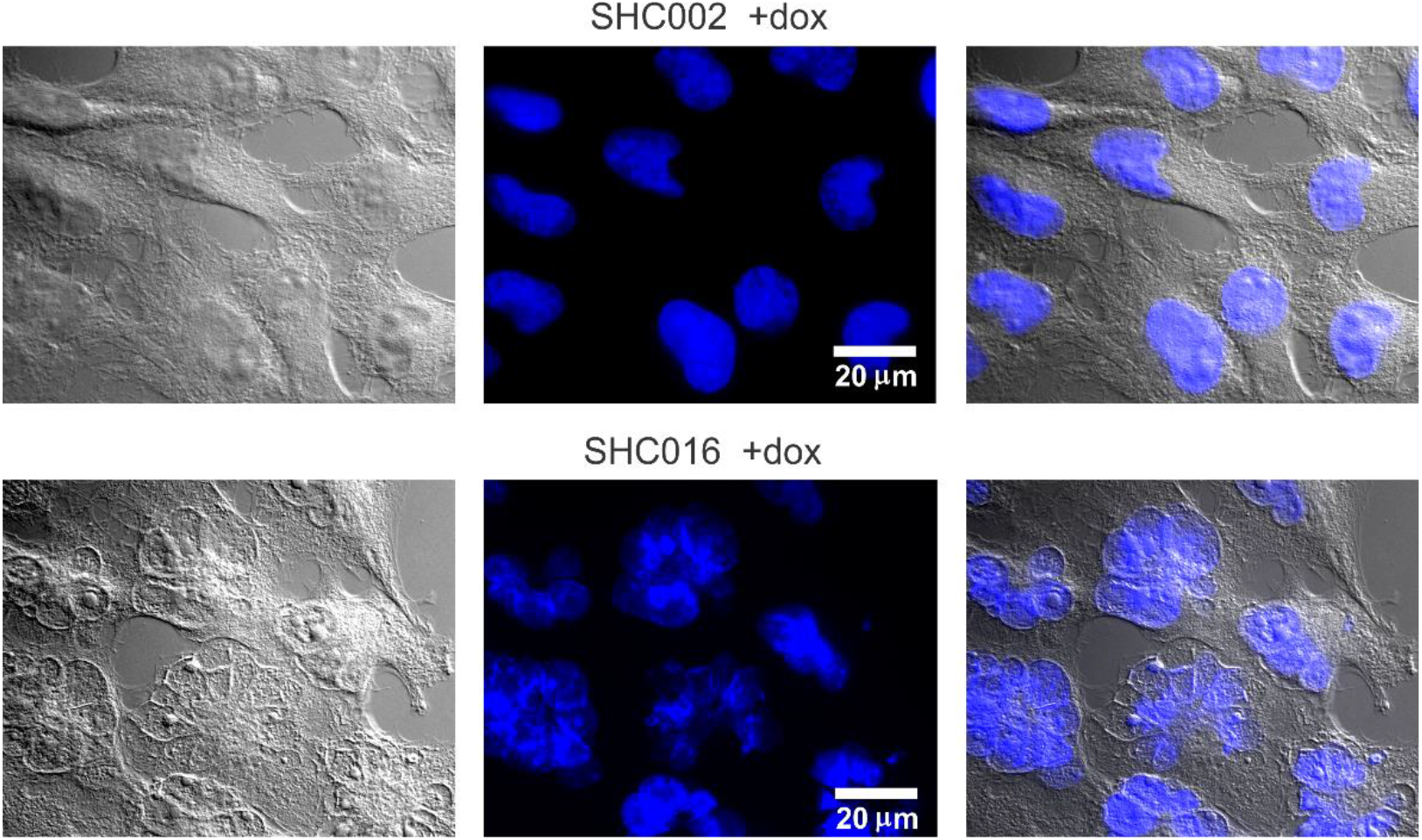
Fluorescence microscopy analyses of nuclear morphology in U251 cells. Expanded version of Figure 4A of the main text. Original transmitted light- and fluorescence images of U251 cells are presented apart from the merged ones. DNA (blue) was stained with DAPI.

**Figure supplement 1B.**
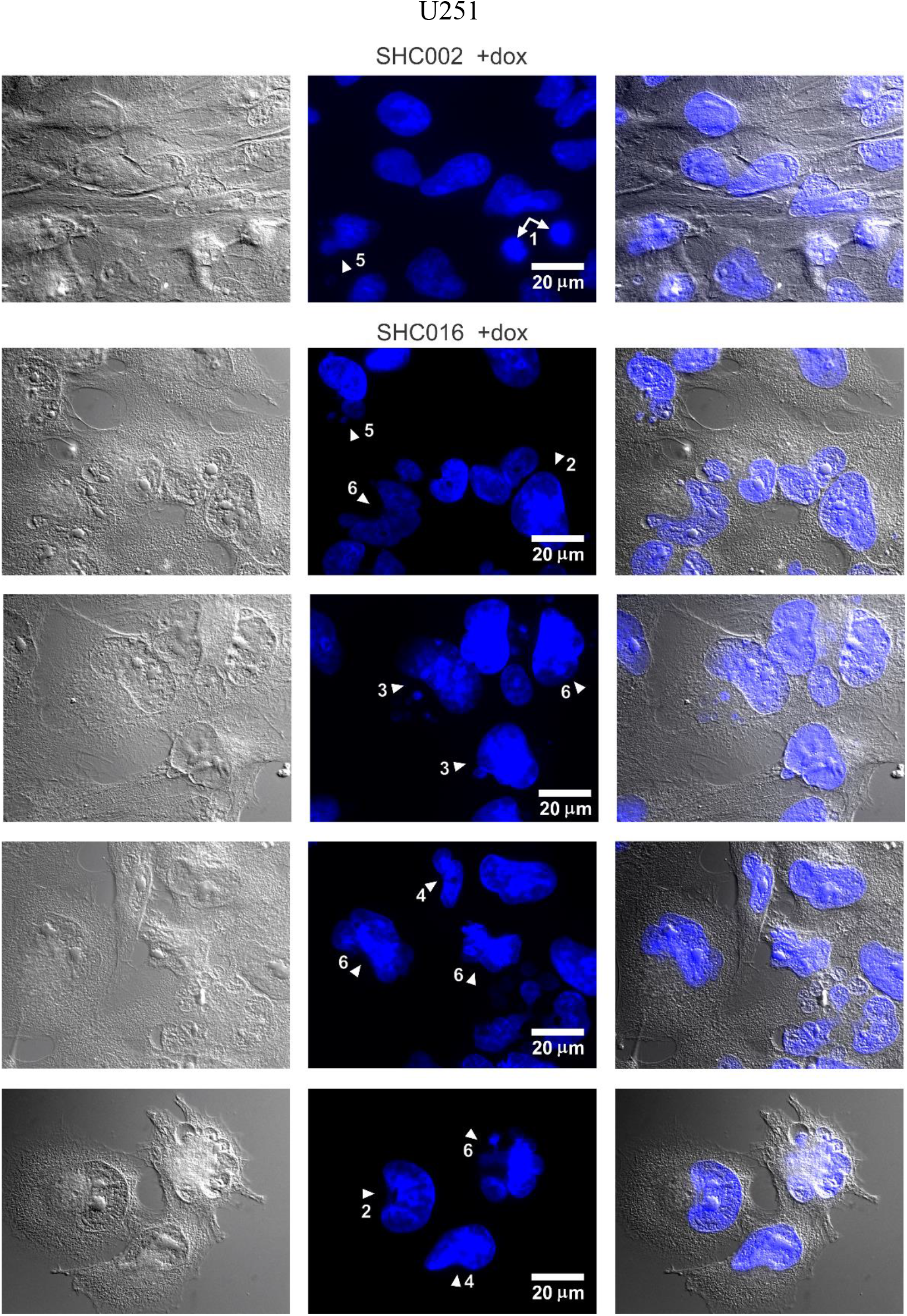
Fluorescence microscopy analysis of nuclear morphology in U251 cells – additional examples. Transmitted light images (left panel), fluorescence images of DNA stained with DAPI (middle panel) and merged images (right panel) of U251 cells taken 5 days after inducing expression of SHC002 or SHC016 shRNAs. Arrowheads point out exemplary features: 1–cytokinesis, 2– enlarged nucleus, 3–enlarged nucleus with micronuclei, 4–nucleus of irregular shape, 5– nucleus of irregular shape with micronuclei, 6–mitotic catastrophe.

**Figure supplement 1C.**
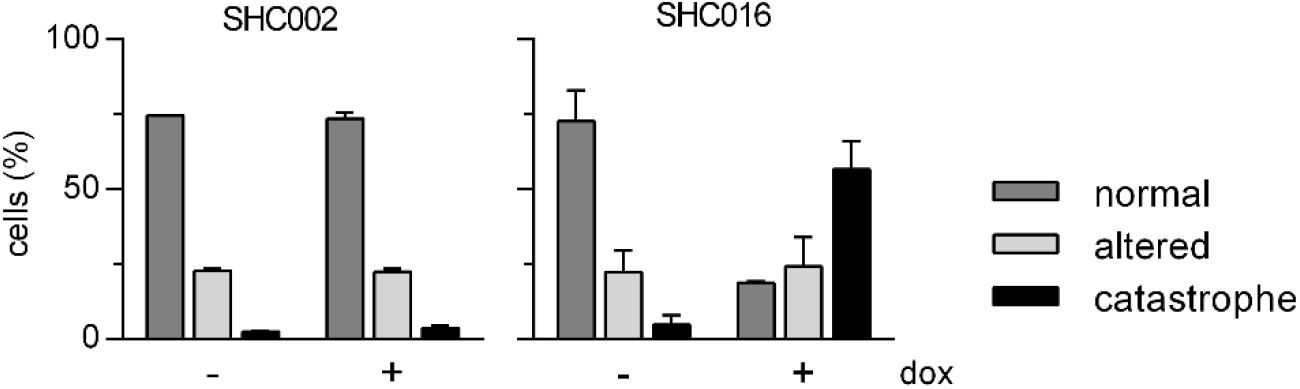
Quantitative analysis of nuclear morphology in U251 with uninduced or induced expression of SHC002 or SHC016 shRNAs – expanded version of Figure 4B of the main text. MV ± SD from two independent experiments are presented; in each experiment at least 120 cells were analyzed in each experimental group. The population described as ‘altered’ includes cells with significantly enlarged nuclei, nuclei with irregular shapes, and those with micronuclei.

**Figure supplement 1D.**
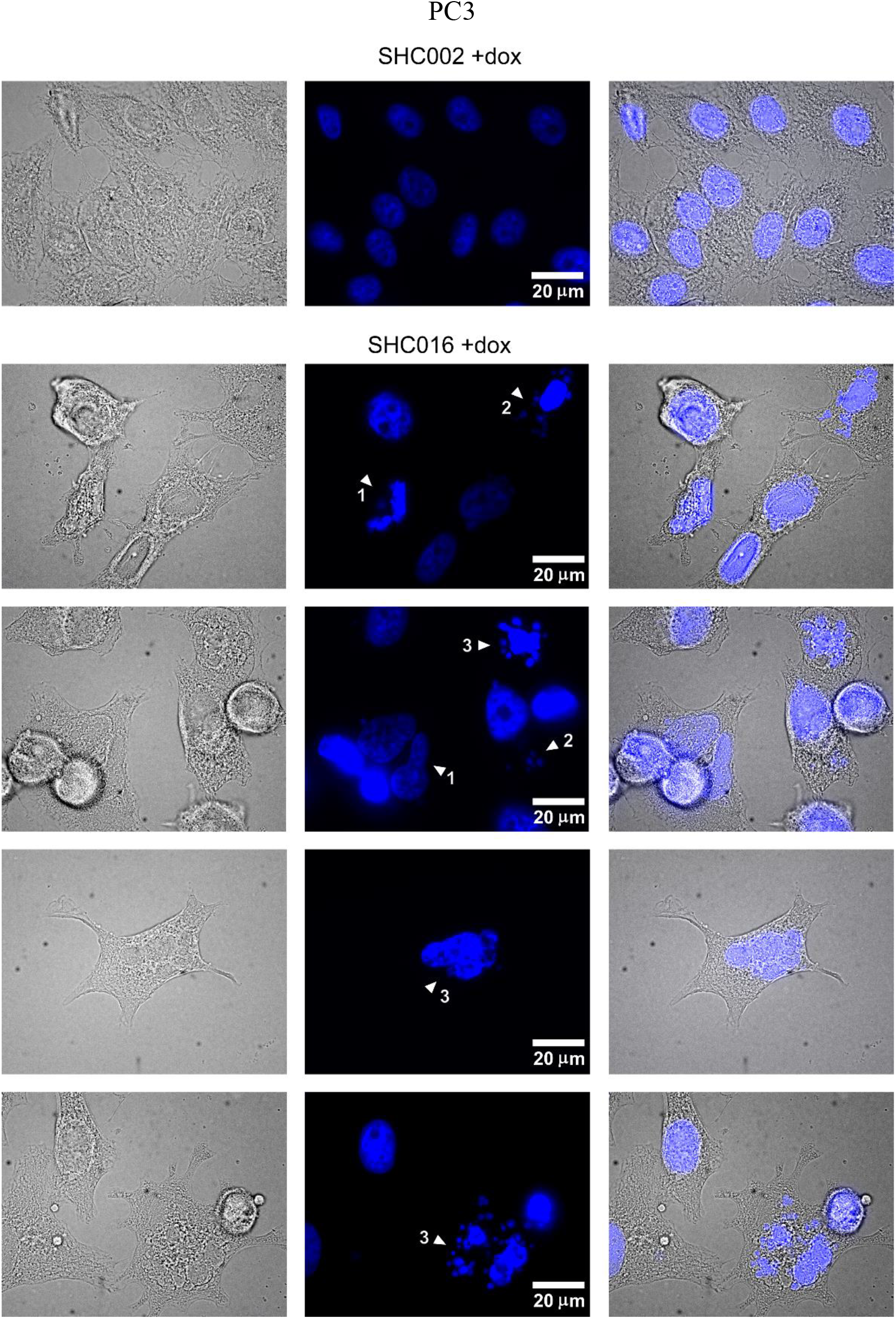
Fluorescence microscopy analysis of nuclear morphology in PC3 cells. Transmitted light images (left panel), fluorescence images of DNA stained with Hoechst 33342 (middle panel) and merged images (right panel) of PC3 cells taken 5 days after inducing expression of SHC002 or SHC016 shRNAs. Arrowheads point out exemplary features: 1–irregular shape, 2– micronuclei, 3–mitotic catastrophe.

**Figure supplement 1E.**
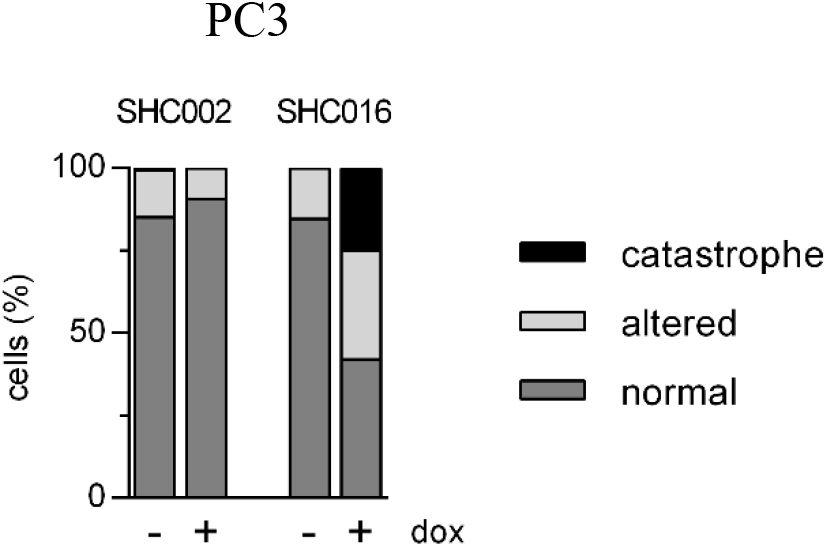
Quantitative analysis of nuclear morphology in PC3 with uninduced or induced expression of SHC002 or SHC016 shRNAs. Data are from a single experiment in which at least 100 cells were analyzed in each experimental group. Similar changes in nuclear morphology of SHC016-expressing PC3 cells were observed in additional experiments. The population described as ‘altered’ includes cells with nuclei of irregular shapes and those with micronuclei.

**Figure supplement 1F.**
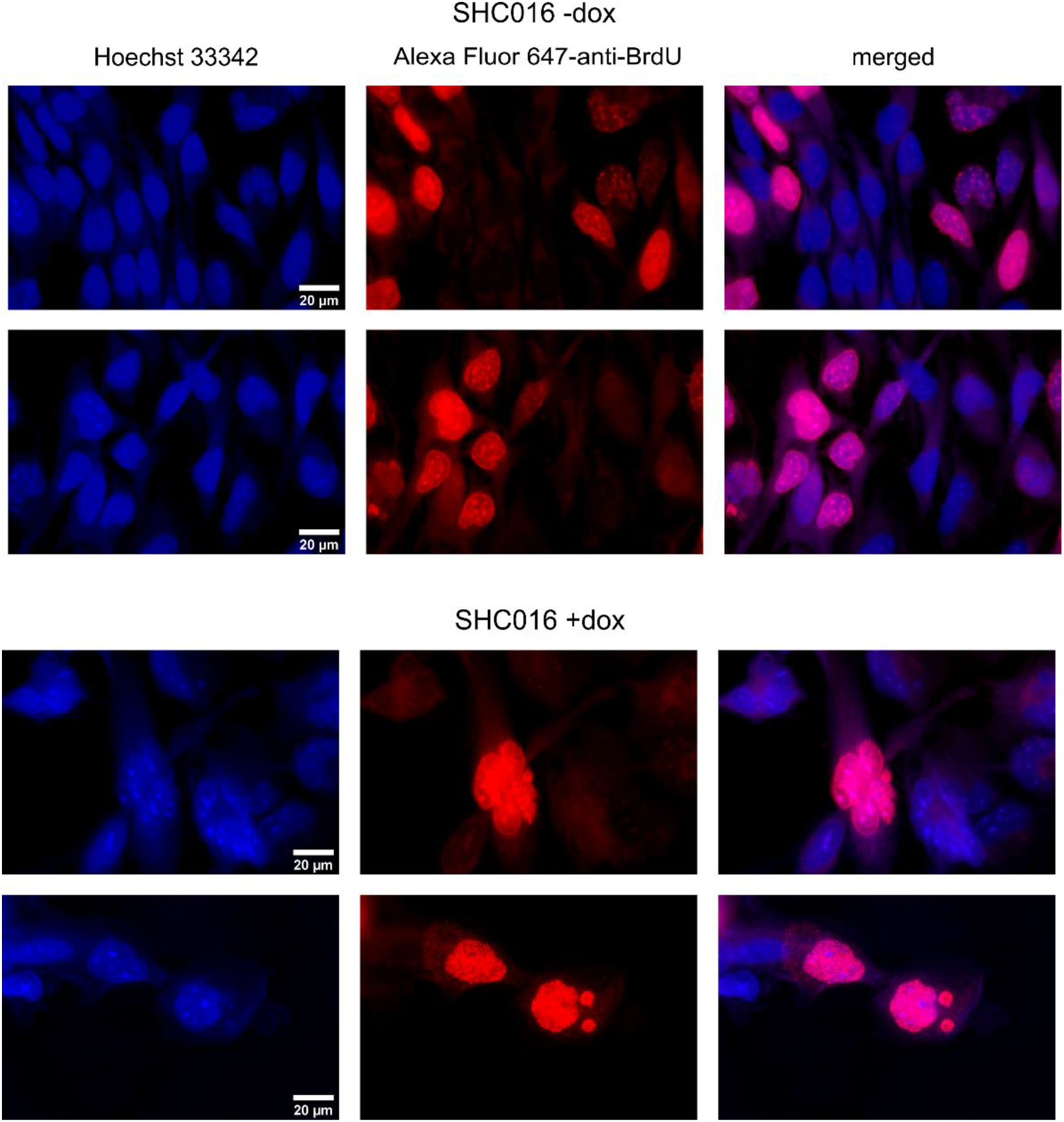
Fluorescence microscopy analysis of BrdU incorporation into DNA of U251 cells, in which the expression of SHC016 was not induced (upper panels) or was induced with doxycycline 5 days prior to staining (lower panels). Similar results were obtained with Alexa Fluor 488-conjugated anti-BrdU antibodies.

**Figure supplement 2.**
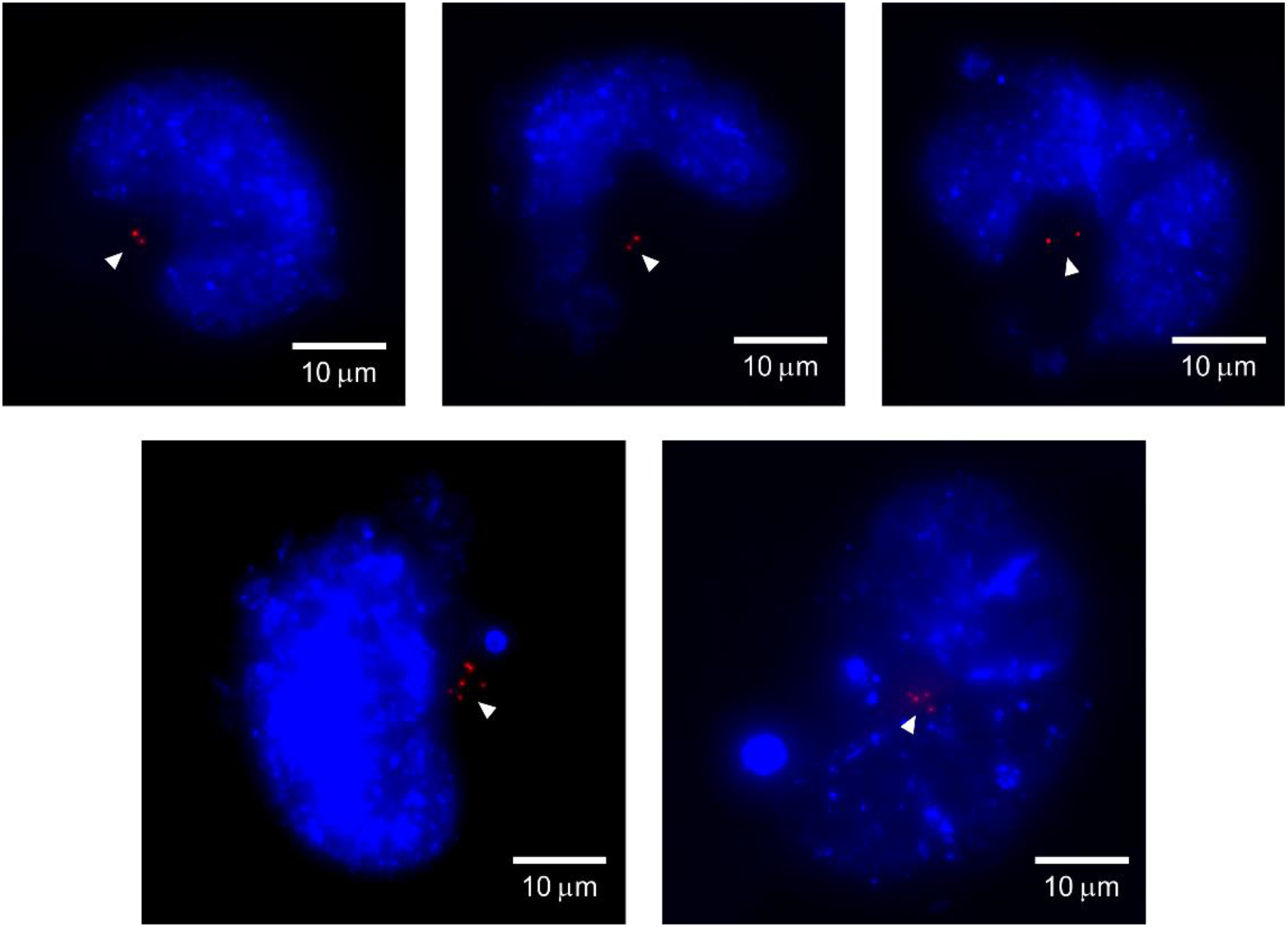
Additional examples of fluorescence images of interphase nuclei of SHC016-expressing cells. DNA was stained with Hoechst 33342 (blue). Arrowheads point out pairs of centrosomes (upper panel) or multiplied centrosomes (lower panel) visualized by γ-tubulin immunostaining (red).

**Figure supplement 3.**
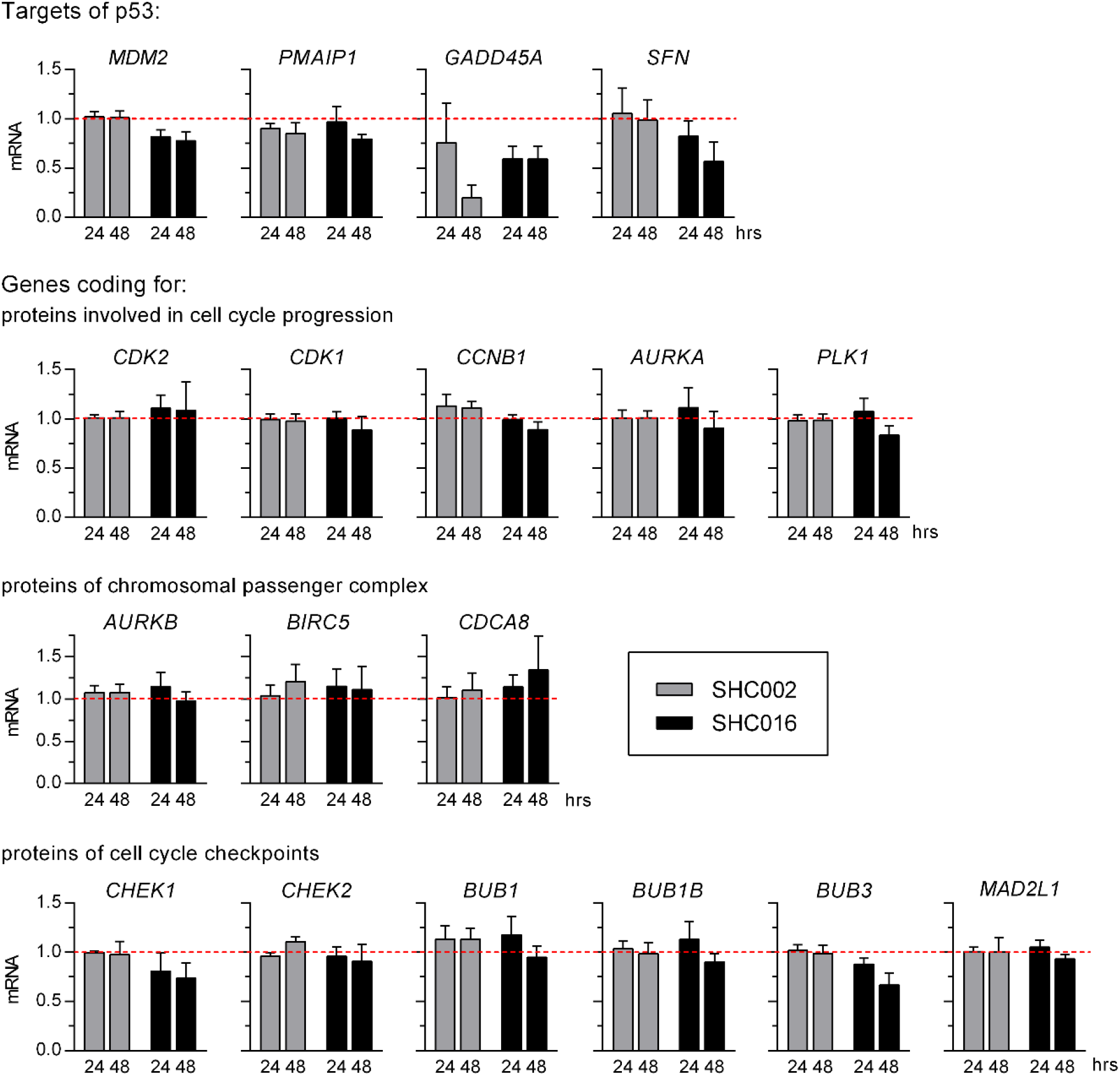
RT-qPCR analysis of the levels of selected transcripts in U251 cells. The analyzed mRNAs comprise targets of p53 as well as those coding for proteins involved in cell cycle progression and control. mRNA was isolated 24 or 48 h after inducing expression of SHC002 or SHC016. The relative levels of the transcripts in uninduced cells were taken as 1. The primers’ sequences are available in *Manuscript supplement—Table I*. MV ± SD from 3 independent experiments are shown.

#### Figure 5

**Figure supplement 1.**
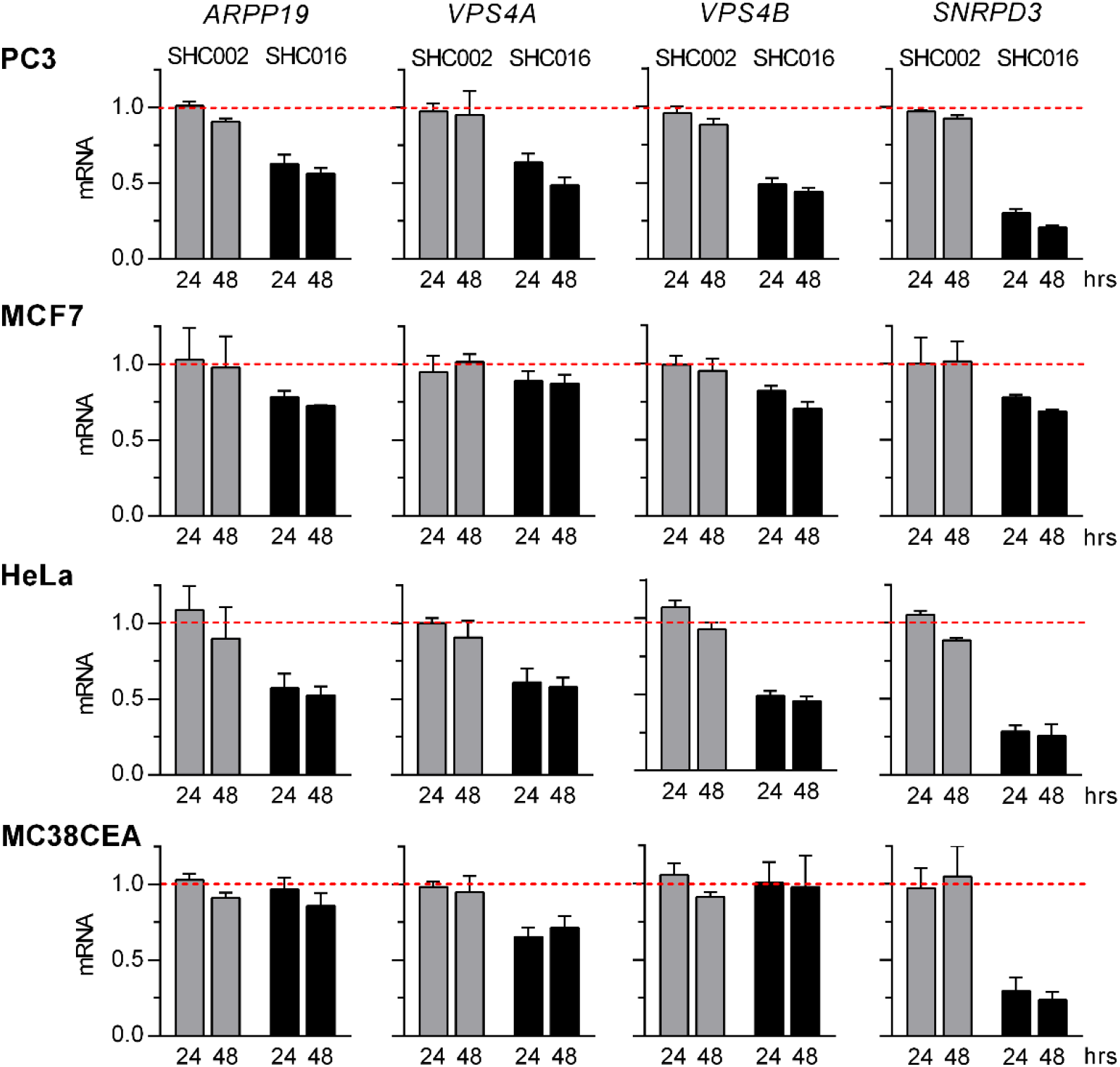
RT-qPCR analysis of transcripts identified by RNAseq as potentially affected by SHC016 shRNA performed 24- and 48 h after inducing expression of SHC002 or SHC016 in human PC3, MCF7, and HeLa cells and in murine MC38CEA cells. The relative levels of the transcripts in uninduced cells were taken as 1. The primers’ sequences are available in *Manuscript supplement—Table I.* MV ± SD of relative mRNA levels from three independent experiments are shown.

**Figure supplement 2.**
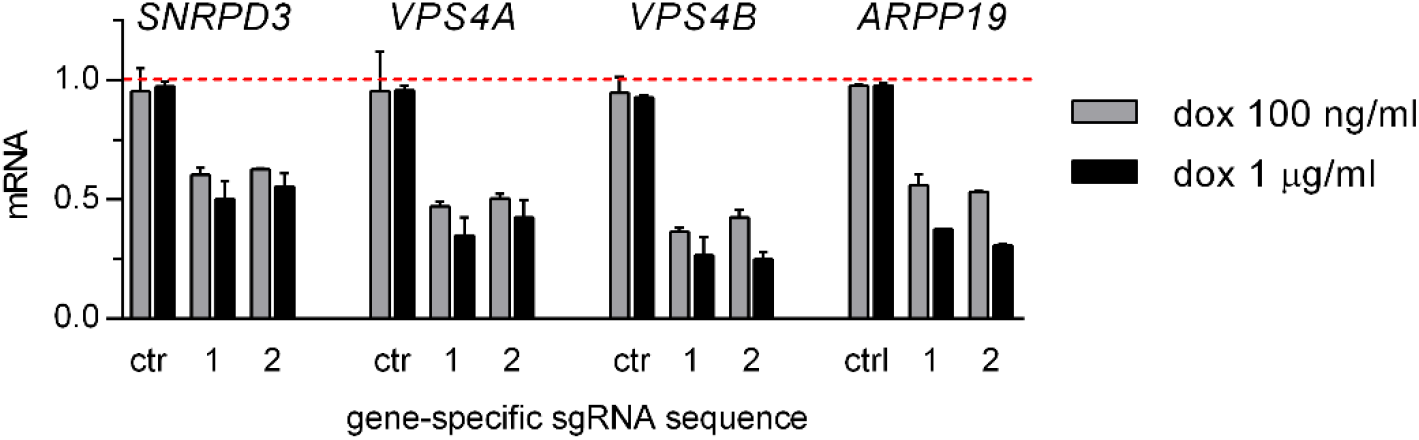
Expanded version of the upper panel of Figure 5B of the main text. RT-qPCR analysis of the extent of gene silencing upon inducing of dCas-KRAB-MeCP2 with doxycycline (dox) used at the concentration of 100 ng/ml or 1 μg/ml. RNA was isolated 72 h after addition of dox to U251 cell cultures. Ctr – control cells transfected with an empty CRISPRi plasmid, which does not contain any sequence coding for specific gene-targeting sgRNA. The relative levels of the transcripts in uninduced cells were taken as 1. MV ± SD of relative mRNA levels from three (dox conc. = 1 μg/ml) or two (dox conc. = 100 ng/ml) independent experiments are shown.

**Figure supplement 3.**
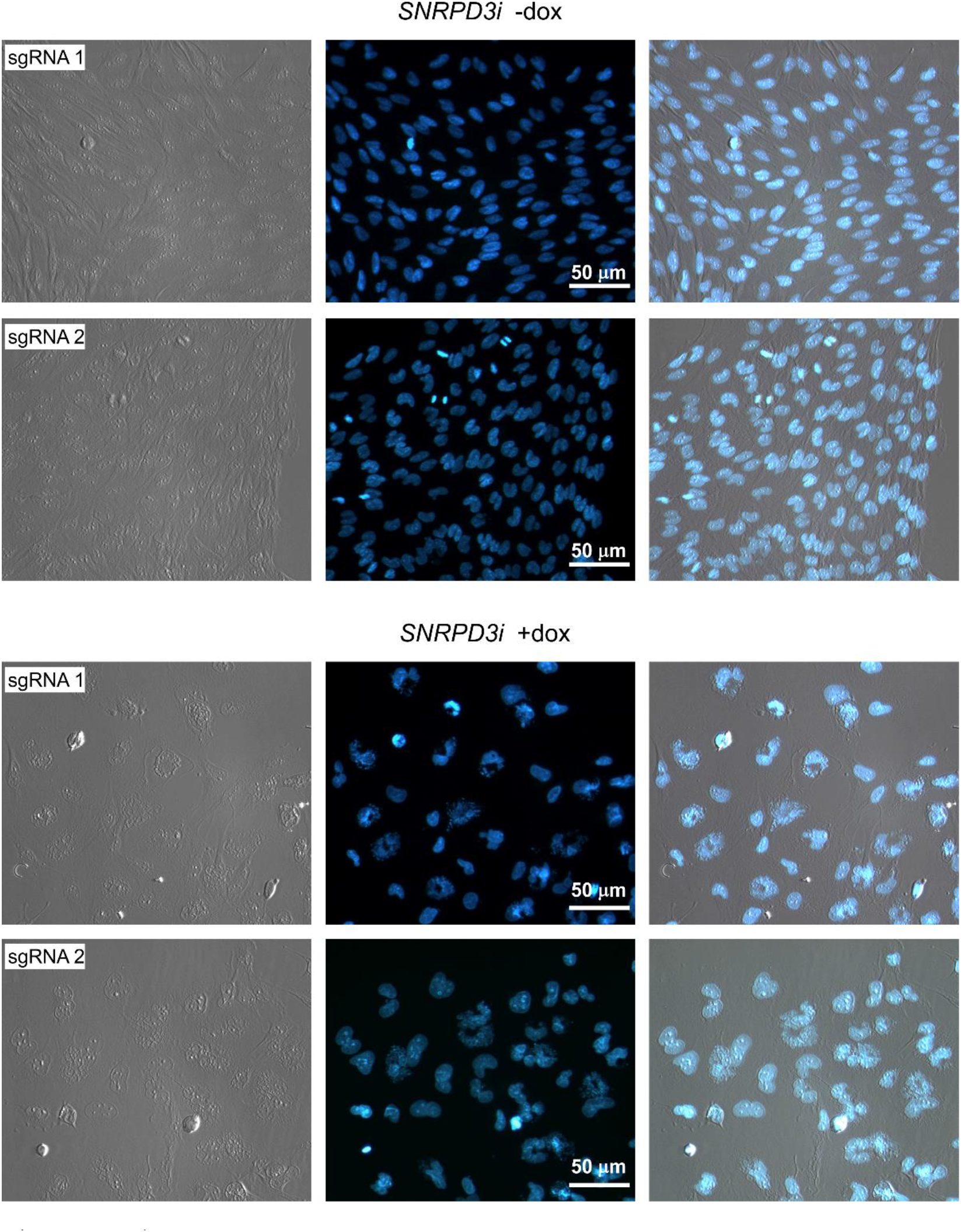
Silencing of *SNRPD3* in U251 cells results in mitotic catastrophe. Fluorescence microscopy analysis of nuclear morphology of U251 cells transfected with *SNRPD3*- targeting inducible CRISPRi vectors, containing either sgRNA 1- or sgRNA 2-coding sequence. Two upper panels present uninduced cells and two lower panels present cells with doxycycline (dox)-induced expression of dCas9-KRAB-MeCP2 six days prior to DNA staining with Hoechst 33342. Left – transmitted light-, middle – fluorescence-, right – merged images.

**Figure supplement 4.**
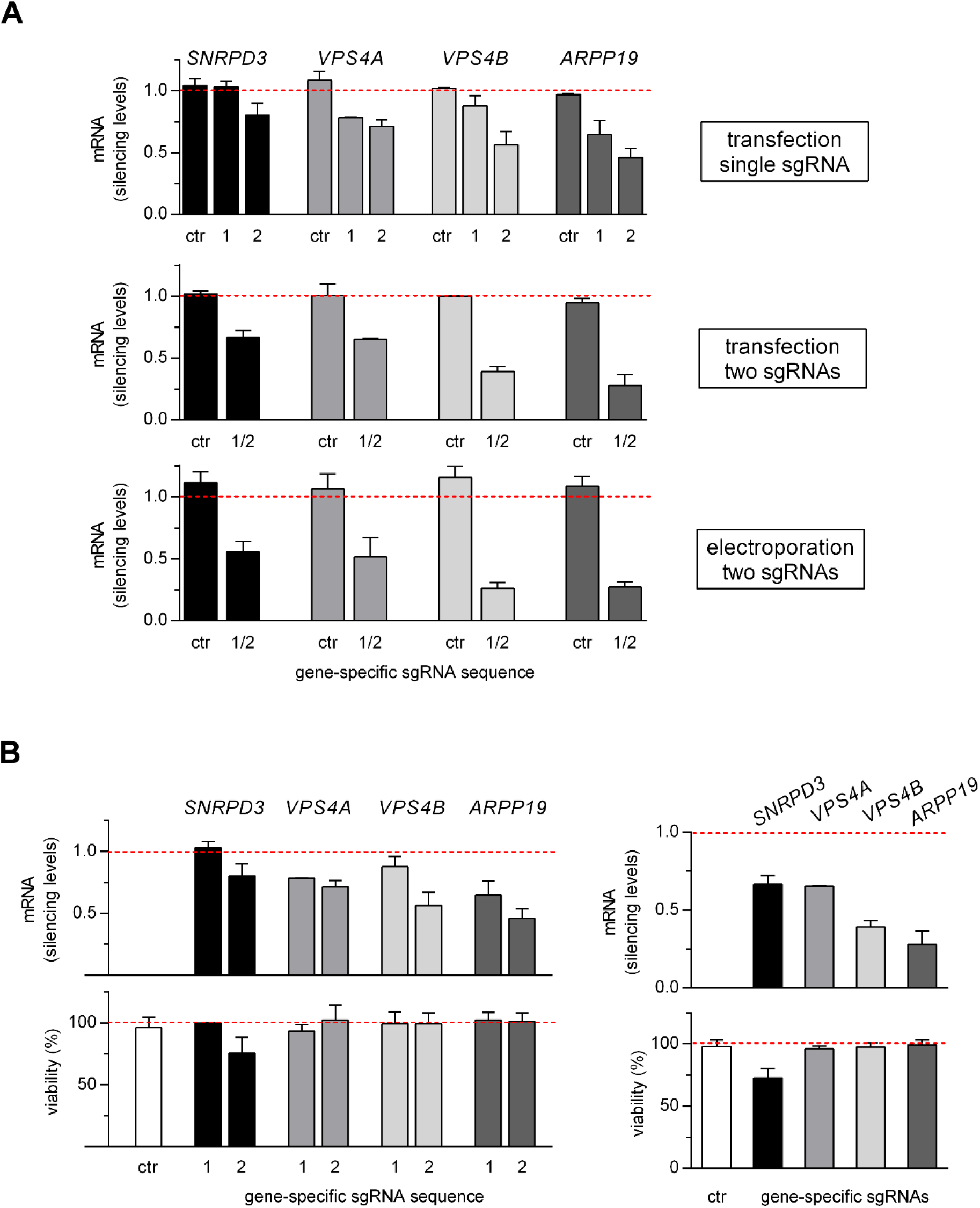
Expanded version of Figure 5C of the main text. (A) Effectiveness of CRISPRi-gene silencing in A549 cells depends on a number of sgRNAs and a method of vector delivery. MV ± SD from two (transfections) or three (electroporation) experiments are presented. (B) Analysis of the correlation between *SNRPD3-* and other genes’ silencing efficiency and viability of A549 cells transfected with CRISPRi vector coding for a single sgRNA, #1 or #2 (left figure) or transfected simultaneously with both CRISPRi vectors encoding two different sgRNAs, #1 and #2 (right figure). Ctr – control cells transfected with an empty CRISPRi plasmid, which does not contain any sequence coding for specific gene-targeting sgRNA. The relative levels of the transcripts in uninduced cells were taken as 1. MV ± SD from two (RT-qPCR) or three (MTT assay) independent experiments are shown.

**Table I.**
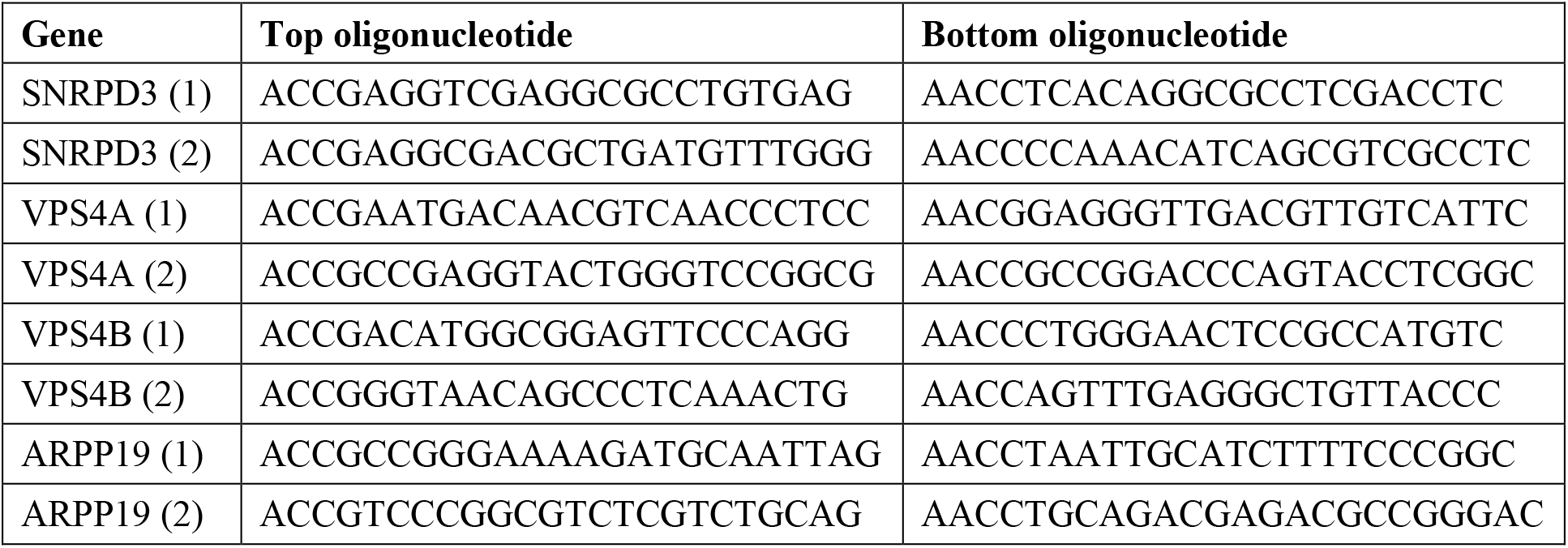
Sequences of sgRNAs

#### Figure 6

**Figure supplement 1.**
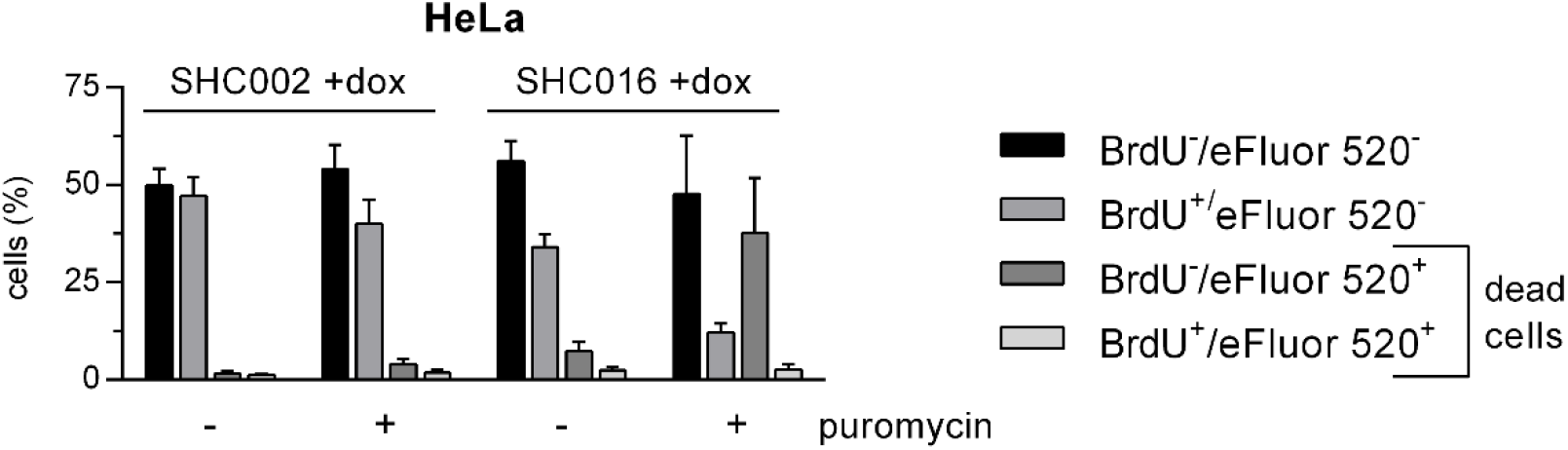
Expanded version of data presented in Figure 6B. DNA synthesis and cell viability assessed via double BrdU/eFluor 520 viability staining of HeLa cells expressing for 5 days either SHC002 or SHC016 in the absence or presence of selection antibiotic, puromycin. Dead – all eFluor 520^+^ cells, BrdU^+^ – eFluor 520^−^/BrdU^+^ cells, BrdU^−^ – eFluor 520^−^/ BrdU^−^ cells. MV ± SD of three independent experiments are shown.

#### Discussion

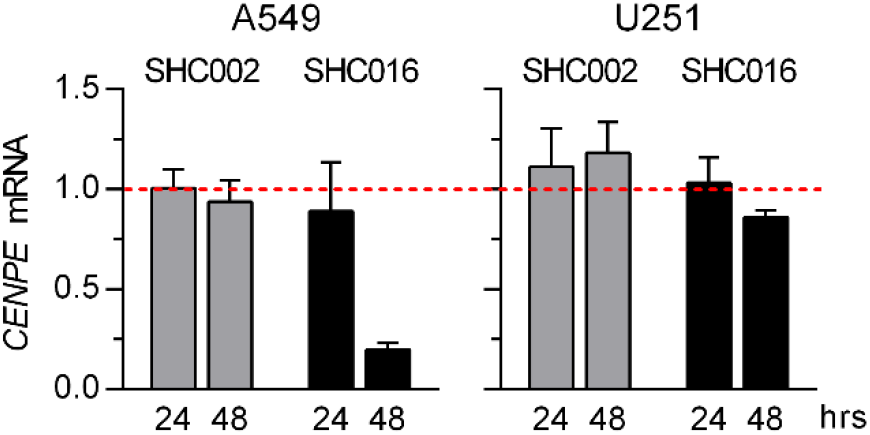

Discussion supplement 1

RT-qPCR analysis of *CENPE* mRNA levels performed 24- and 48 h after inducing expression of SHC002 or SHC016 in A549 and U251 cells. The relative levels of *CENPE* transcript in uninduced cells were taken as 1. MV ± SD of relative mRNA levels from three independent experiments are shown.

Manuscript supplement—Table I

**Table I.**
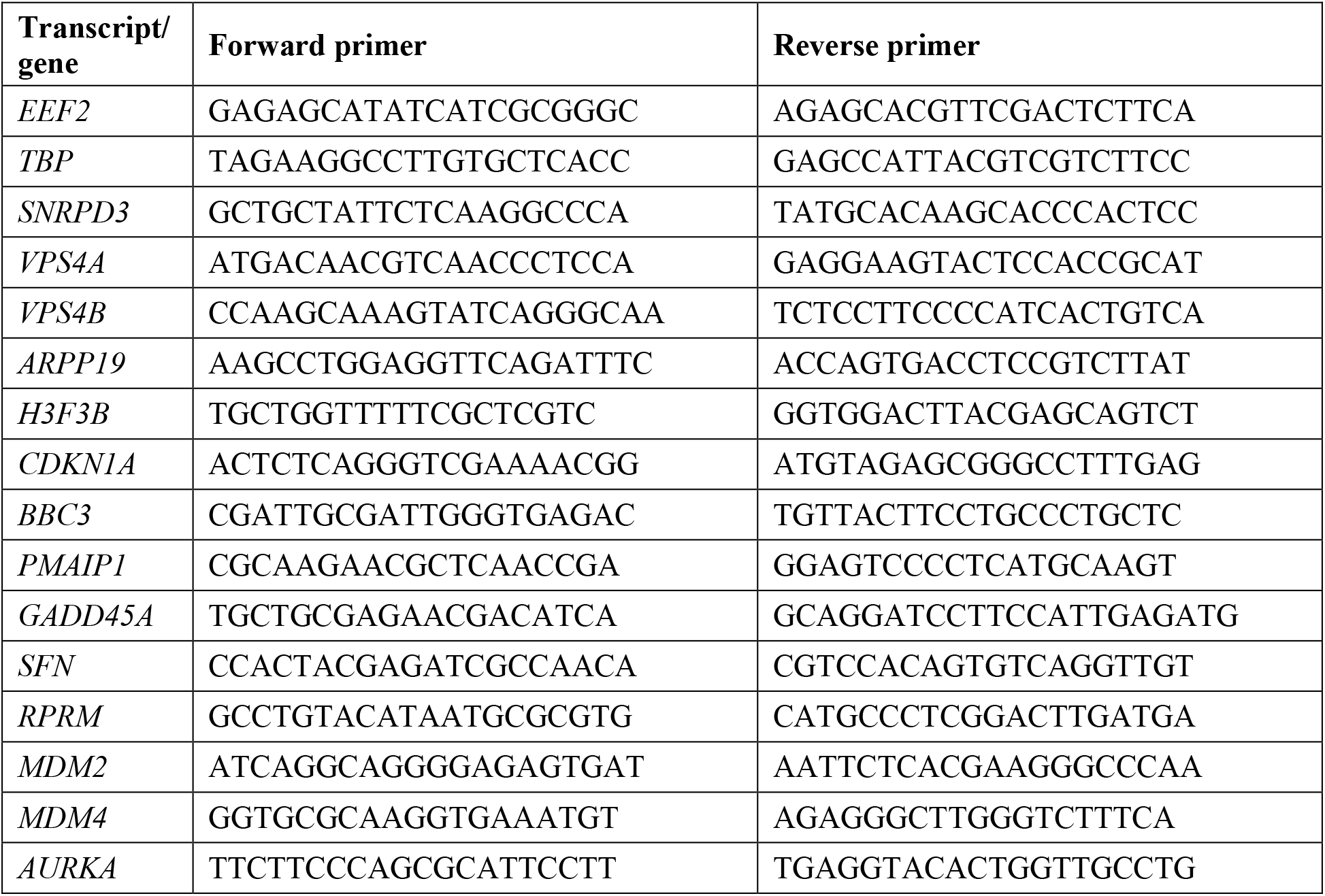
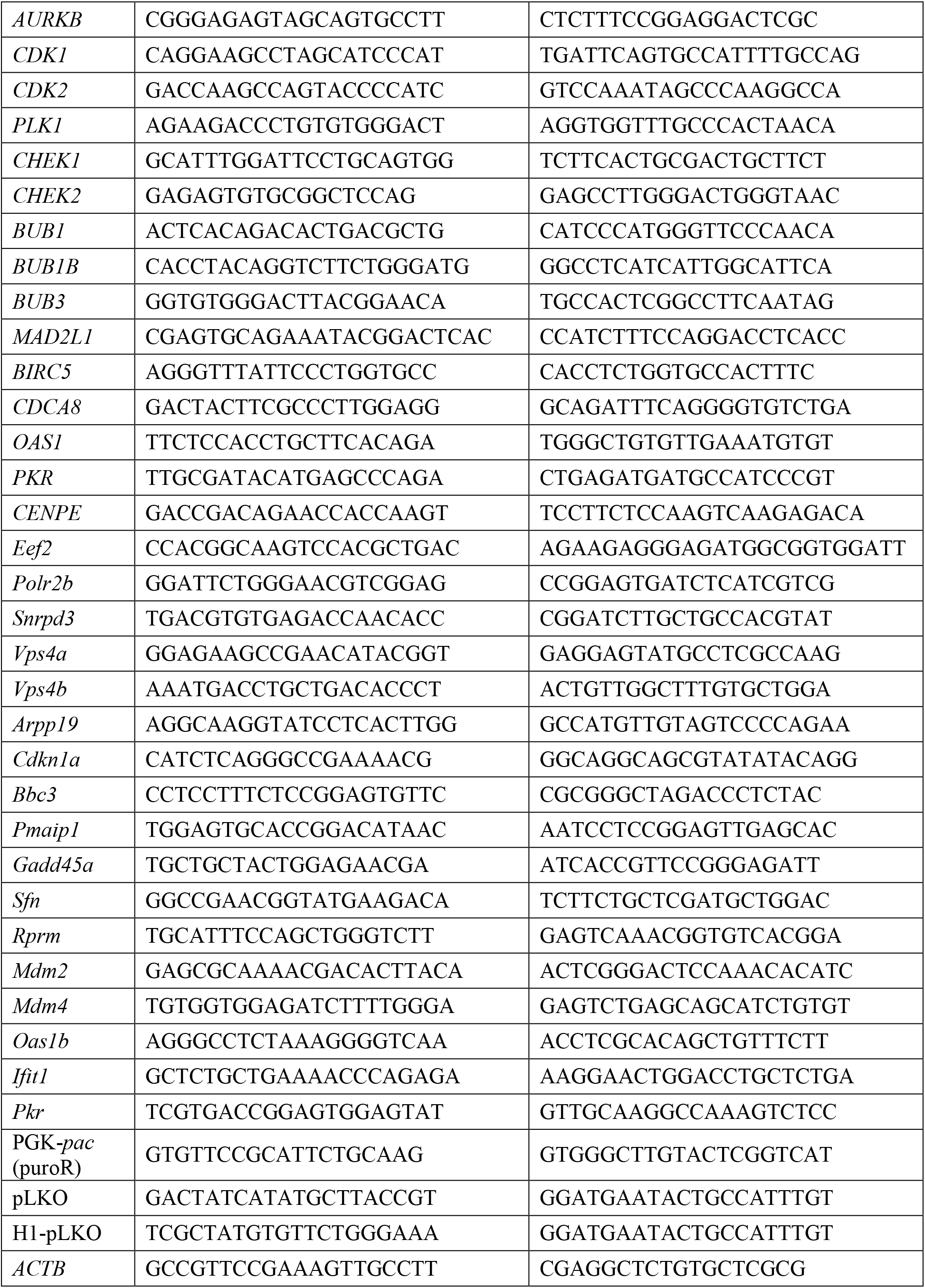
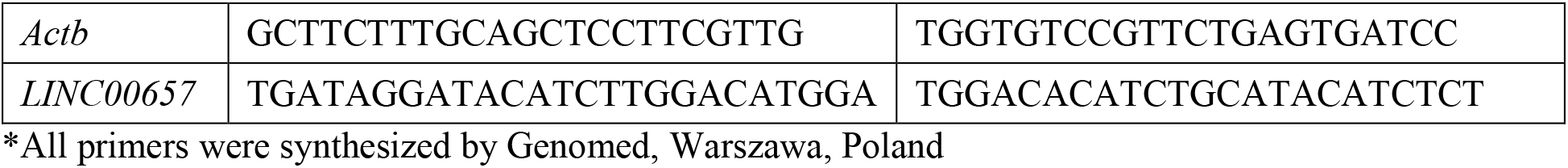
Sequences of primers*

